# An integrated mass spectrometry strategy for quantifying the proteoform diversity of the extensively modified O-glycoprotein Osteopontin

**DOI:** 10.64898/2026.09.07.749945

**Authors:** Konstantin C. Zouboulis, Jack L. Bennett, Leonard A. Daly, Sean A. Burnap, Ellie Holden, Corinne A. Lutomski, Claire E. Eyers, Carol V. Robinson, Weston B. Struwe, Justin L.P. Benesch

## Abstract

O-glycosylation is among the most abundant and structurally diverse post-translational modifications in eukaryotes, yet its heterogeneity renders O-glycoproteins exceptionally difficult to characterize. To overcome these challenges, we have developed an integrated mass spectrometry (MS) strategy to define the proteoform landscape of O-glycoproteins and applied it to human osteopontin (OPN). OPN is a disease-associated extracellular matrix protein subject to extensive modification. By combining native MS with serial exoglycosidase digestions, we directly resolved truncation, phosphorylation, sulfation, and O-glycosylation of OPN. Matched glycoproteomic analyses, using tailored (glyco)protease combinations, allowed us to quantify glycan heterogeneity inaccessible to conventional trypsin-based approaches or protein-centric methods. We integrated these datasets using forward compositional simulations to infer the intact OPN proteoform distribution and benchmarked the resulting models against an experimental intact-mass distribution obtained by proton-transfer charge-reduction MS. This comparison revealed that bottom-up O-glycoproteomics systematically underestimates the true extent of glycan sialylation, whereas assuming (near-)complete sialylation accurately reproduced the experimental intact OPN mass distribution. Together, these results provide a comprehensive, quantitative view of OPN compositional diversity and demonstrate how intact-protein and peptide-level measurements can be reconciled to resolve highly heterogeneous glycoform populations. The workflow establishes a broadly applicable framework for characterizing extensively O-glycosylated and multiply modified proteins.

## Introduction

O-glycosylation is among the most abundant and structurally diverse post-translational modifications (PTMs) in eukaryotes, governing protein folding, stability, and intermolecular recognition.^1–3^ Unlike N-glycosylation, O-glycosylation lacks a consensus sequon but is similar in possessing two compounding layers of structural complexity: macroheterogeneity, the variable occupancy of a given site across the proteoform population, and microheterogeneity, the range of glycan structures present at any single occupied site.^4,5^ When multiple O-glycan sites are present, these sources of heterogeneity combine to generate large and highly complex proteoform populations.

Peptide-centric, bottom-up glycoproteomics has established itself as the method of choice for glycoprotein analysis, achieving glycosite- and glycan composition-level resolution.^4,6–8^ However, it is now increasingly recognized that peptide-centric analysis alone cannot provide a complete description of the overall protein glycosylation state.^9–11^ Protein-centric mass spectrometry (MS) approaches provide a complementary view, capturing the proteoform population as a whole and preserving the relationships between co-occurring modifications.^9,12,13^ The application of protein-centric approaches, however, has been largely restricted to comparatively simple glycosylated systems.^9,10,14–16^ In extensively glycosylated proteins, the number of overlapping compositions can produce severe spectral congestion, preventing direct assignment of individual proteoforms.

Recent measurement strategies have begun to address this limitation.^17^ Charge- detection MS^11^ and proton-transfer charge-reduction MS (PTCR MS)^18^, for example, can recover accurate intact-mass distributions from highly heterogeneous samples. However, individual molecular features still often remain masked by glycan heterogeneity, and can only be observed through enzymatic glycan release^5,19^ or top- down fragmentation^20,21^. Thus, no single MS measurement provides the level of compositional insight necessary to completely describe an extensively O-glycosylated protein. Integrated workflows that deliberately combine protein- and peptide-centric MS modalities are therefore required to resolve glycoform populations.

A prototypical multiply modified O-glycoprotein, osteopontin (OPN), also known as secreted phosphoprotein 1 (SPP1), is an intrinsically disordered matricellular protein that carries extensive O-glycosylation alongside multiple co-occurring PTMs.^22–25^ Widely expressed across osteoblasts, epithelial cells, and immune cells, OPN contributes to physiological processes such as bone remodeling, angiogenesis, and immune modulation through interactions with integrin receptors and CD44 splice variants.^26–31^ OPN overexpression in pathological states has positioned it as a potential disease biomarker and therapeutic target.^32,33^ Notably, OPN is upregulated across a broad range of tumor types, where elevated expression has been associated with poor prognosis and reduced survival.^23,34–37^ In these pathological environments, OPN interactions activate signaling pathways that promote tumorigenesis, tumor growth, and metastasis.^23,35,36,38^

There is growing evidence that OPN’s receptor engagement and signaling are regulated by PTMs acquired during protein maturation, including proteolytic cleavage, phosphorylation, and N-/O-glycosylation (Figure 1A).^36,39–42^ N-glycosylation is inconsistently reported and appears to depend on tissue origin and study, despite two potential N-X-S/T consensus sequons.^22,43–45^ O-glycosylation, in contrast, is consistently observed across cell and tissue types.^22,41,44–47^ Edman degradation and matrix-assisted laser desorption/ionization MS (MALDI-MS) identified five O- glycosylated threonine residues (Thr134, Thr138, Thr143, Thr147, Thr152) within a threonine/proline-rich region of milk- and urine-derived OPN.^22,44^ Subsequent tandem MS studies on HEK293-derived OPN provided only partial support for these sites, while also reporting additional glycosites (Figures 1B, S1).^41,45,46^ Functionally, sialylation of O-glycans influences integrin binding,^39^ and mutating O-glycosylation sites enhances cell adhesion and spreading, an effect further increased upon dephosphorylation.^41^ Mutations at Thr143, Thr147, and Thr152, located near the conserved RGD^159–161^ integrin-binding motif, increase OPN-mediated adhesion in cancer cell lines. OPN is also extensively phosphorylated, with more than 36 sites reported across the protein.^22,45,48–51^ C-terminal phosphorylation reduces OPN-mediated integrin-dependent adhesion relative to less phosphorylated proteoforms.^40^ Phosphorylation may also be influenced by the extent of O-glycosylation, suggesting biosynthetic crosstalk between these PTMs.^41,46^ Resolving how these modifications coexist across the OPN population is therefore essential for understanding how proteoform diversity shapes its physiological and pathological functions.

**Figure 1:**
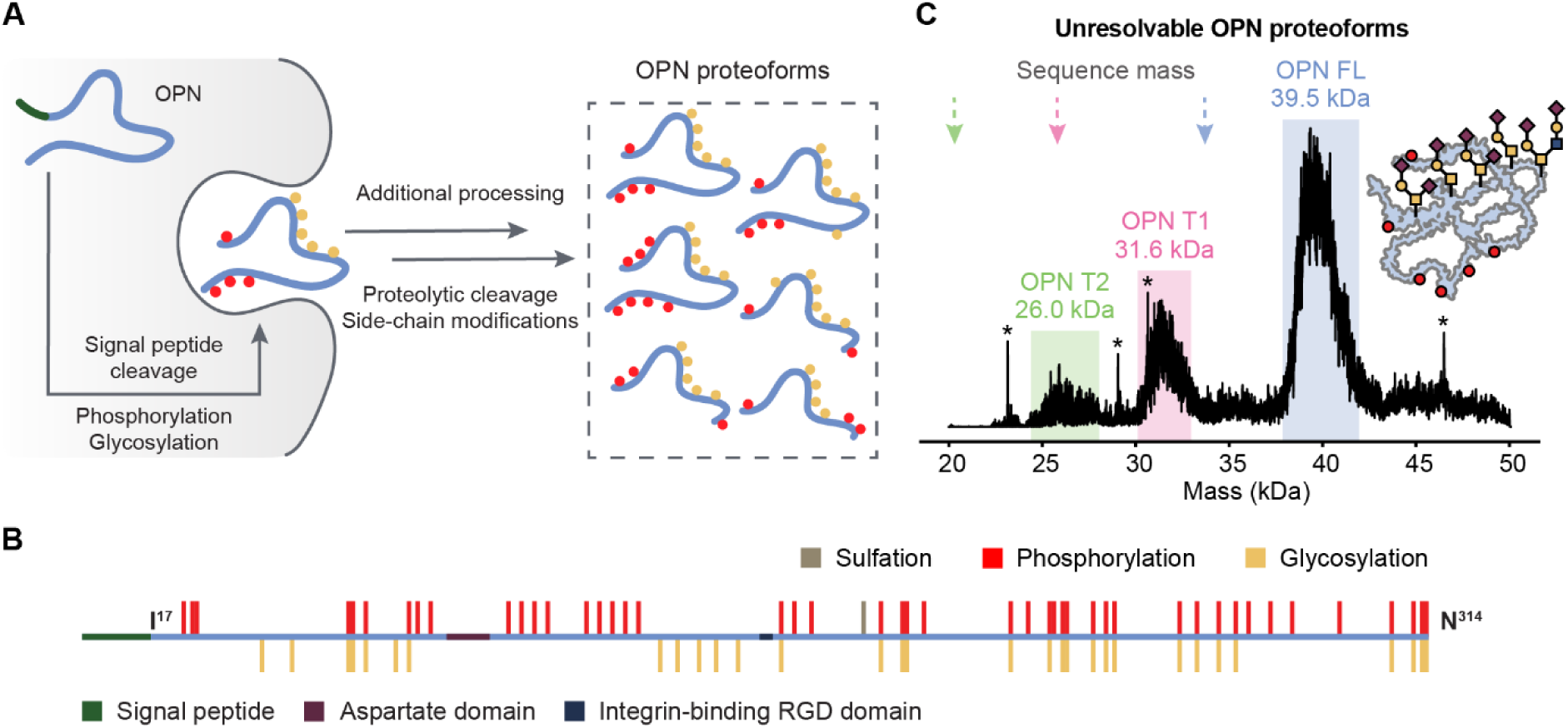
**(A)** During secretion and maturation, OPN undergoes signal peptide (SP) cleavage and extensive post-translational modification (PTM), generating a diverse set of proteoforms. **(B)** Schematic representation of the OPN sequence with selected sequence features highlighted including the N-terminal SP, the aspartate domain DDMDDEDDDD^86–95^ and the RGD^159-161^ integrin binding motif. PTM modifications reported in the literature^22, 41,44,45,48,51^ include sulfation, phosphorylation, and O- glycosylation. **(C)** The deconvolved DIA-PTCR MS/MS mass spectrum of OPN contains three broad peaks corresponding to full-length OPN (FL) and its truncated forms (T1 and T2). Arrows mark the sequence-predicted masses of 33.7 kDa, 25.8 kDa, and 20.3 kDa for OPN FL, OPN T1, and OPN T2, respectively.

To enable comprehensive analysis of heterogeneously O-glycosylated proteins such as OPN, here we develop an integrated, multiscale MS strategy that combines protein- and peptide-centric approaches to uncover glycoform landscapes. Native MS (nMS) with sequential exoglycosidase treatment reveals truncations and overall glycosylation and phosphorylation states directly on the intact protein, while tailored glycoproteomic workflows resolve glycosylation sites and glycan types beyond the reach of conventional trypsin-based approaches. We integrate these measurements through forward compositional simulations and benchmark the reconstructed proteoform distributions against experimental intact-mass distributions obtained by PTCR MS. Our results reveal the extent to which peptide-level measurements capture, and systematically underestimate, features of the intact glycoform population. More broadly, this approach provides a framework for resolving the compositional diversity of extensively O-glycosylated and multiply modified proteins.

## Results and Discussion

### Diverse OPN modifications give rise to a complex array of mass-disperse proteoforms

To accurately characterize human OPN proteoform diversity, we performed nMS experiments on full-length human OPN (OPNa, UniProt P10451-1) expressed in HEK293 cells. OPNa consists of 314 amino acid (aa) residues, including the 16-aa canonical N-terminal signal sequence, which is cleaved during secretion from the cell. The native mass spectrum (Figure S2A) contained broad, unresolved peaks, preventing the resolution of individual proteoforms. Such broad peaks in nMS are often indicative of compositional heterogeneity, which we attributed to variable O-glycosylation, consistent with previous studies reporting O-glycosylation of OPN.^22,40,41,44,45^

To address this spectral complexity, we leveraged ion–ion reactions to differentially reduce the charge state of analyte ions within narrow *m*/*z* isolation windows. Specifically, we employed a data-independent (DIA) PTCR MS/MS strategy, in which narrow *m*/*z* windows (*m*/*z* 15) are sequentially isolated, such that only a subset of ions undergoes charge reduction at a time.^18^ By sliding the isolation window over the *m*/*z* range of 2,845 – 4,415, we generated a series of PTCR MS^2^ spectra that were individually deconvolved into the mass domain (Figure S2). The resulting molecular mass distributions were summed to yield a combined mass spectrum, revealing the molecular mass distribution of the OPN proteoforms. We observed three broad peaks around 39.5 kDa, 31.6 kDa, and 26.0 kDa, which we assigned to full-length OPN (OPN FL) and two truncated forms, OPN T1 and OPN T2, respectively (Figure 1C). The measured mass for OPN FL agrees well with literature values for HEK293 cell-derived OPN of 38.5 kDa^40^, 39.8 kDa^40^, and 39.7 kDa^41^ determined by MALDI-MS. Consistent with this, our previous native top-down MS (nTDMS) analysis of HEK293-derived OPN identified OPN FL as the most abundant proteoform, alongside two C-terminal truncation variants at Tyr246 (OPN T1) and Glu197 (OPN T2).^21^ OPN T1 has also been identified in urine.^44^

Comparing the PTCR mass distribution with the sequence-predicted masses of 33.7 kDa, 25.8 kDa, and 20.3 kDa for OPN FL, OPN T1, and OPN T2, respectively, revealed that the observed masses are substantially higher. The resulting mass differences of approximately 5.8 kDa, 5.8 kDa, and 5.7 kDa are consistent across all three proteoforms. This consistency, together with the magnitude of the shift, points to glycosylation as the dominant contributor to OPN’s mass heterogeneity at the intact level and further suggests that most glycosylation sites are located N-terminal to the C-terminal truncation sites.

### Sequential enzymatic treatment reveals the glycosylation state of intact OPN

We next investigated whether selective removal of OPN PTMs using glycosidases and phosphatases could further define the glycosylation and phosphorylation state of OPN. As an initial assessment, the effects of glycosidase treatment were evaluated by SDS– PAGE. Analysis of untreated OPN revealed three distinct bands (Figure S3A), consistent with OPN FL and two truncation variants identified in the DIA–PTCR MS experiment. OPN treatment with neuraminidase A resulted in an SDS-PAGE gel shift compared to untreated OPN, suggesting that the OPN glycans contain terminal sialic acids. Treatment with a mixture of PNGase F, *O*-glycosidase, neuraminidase A, β1-4 galactosidase, and β-*N*-acetylhexosaminidasef, which together enable removal of N- and most O-glycans, resulted in a further gel shift relative to neuraminidase A treatment alone, consistent with the removal of additional glycans. In contrast, treatment with PNGase F, which exclusively removes N-glycans, produced no detectable gel shift (Figure S3B). This supports the observation that OPN from HEK293 cells is not N-glycosylated despite carrying two N-X-S/T sequons. Finally, we incubated OPN with lambda protein phosphatase (LPP), which has activity toward phosphorylated serine, threonine, and tyrosine residues. No significant gel shift was detected, consistent with no or low levels of phosphorylation. Together, these results indicate that this preparation of OPN carries predominantly sialylated O-glycans and is not extensively phosphorylated.

While SDS-PAGE confirmed that glycosidase treatment altered OPN’s apparent molecular weight, it lacked the resolution to define the underlying proteoforms and glycan compositions (Figure S4). We therefore analyzed glycosidase-treated OPN by nMS, applying glycan-specific exoglycosidases sequentially to reduce glycan heterogeneity, simplify the native mass spectra, and resolve the O-glycan types.^5,9,19^

Initial nMS experiments using neuraminidase A and *O*-glycosidase, which together remove sialylated core-1 and core-3 O-glycans, did not yield readily interpretable spectra. We therefore hypothesized that OPN carries extended and/or branched O-glycan structures, such as core-2, that are not removed by these two enzymes. In contrast to Chinese hamster ovary (CHO) cells, which have served as the source for many previous MS-based glycosylation studies^9,18,19^ but have a restricted repertoire of endogenous O-glycan structures and produce predominantly core-1 O-glycans,^52,53^ human HEK293 cells exhibit a more complex O-glycosylation pattern, synthesizing both core-1 and core-2 O-glycans (Figure 2A).^53–55^ To selectively remove these glycans and generate simplified mass spectra, we selected three additional glycosidase combinations (Figure 2B, C).

**Figure 2:**
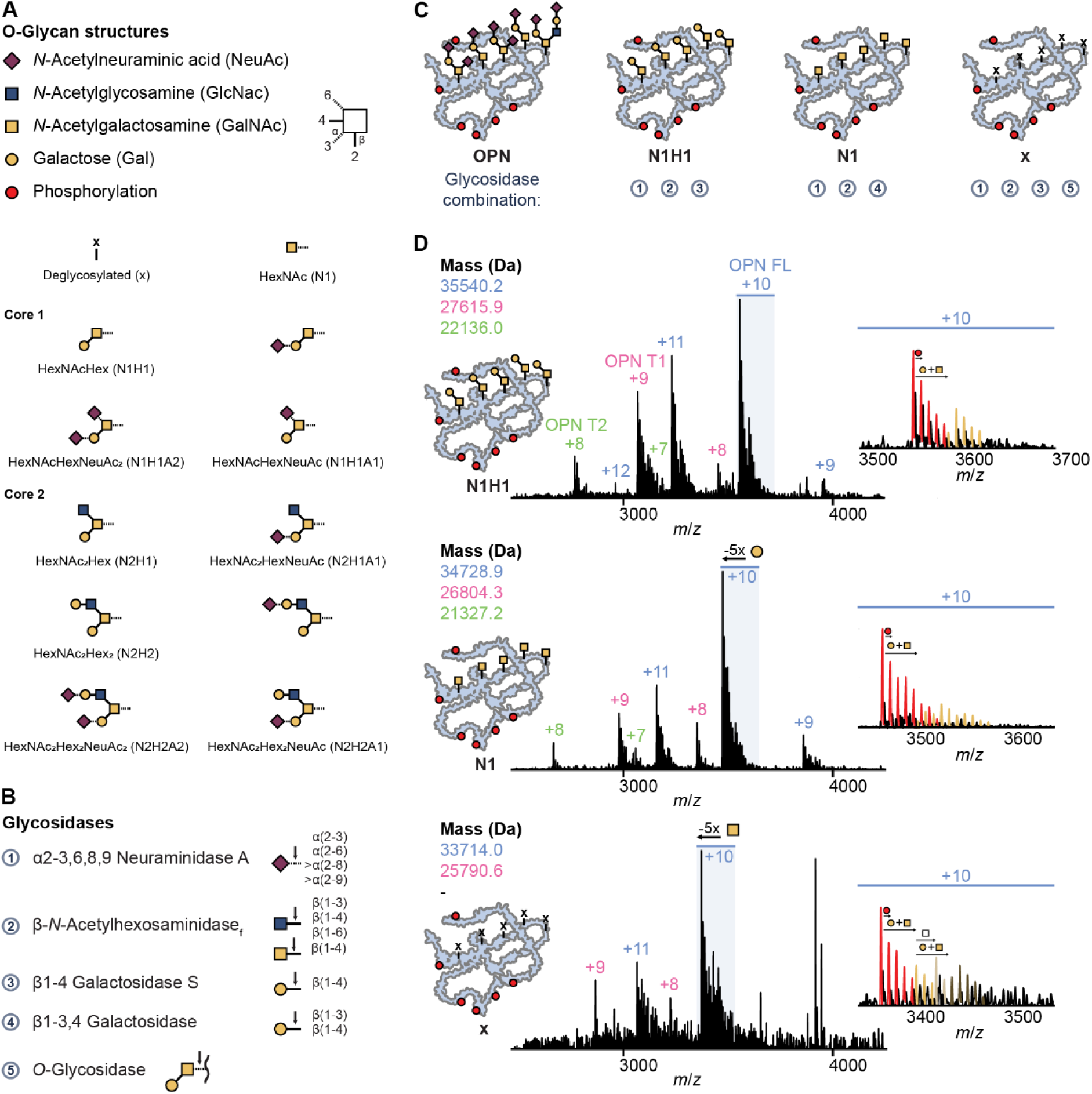
**(A)** Common core-1 and core-2 O-glycan structures. **(B)** Glycosidases and their substrate specificity. **(C)** Controlled enzymatic glycan trimming is hypothesized to remove heterogeneity and generate distinct OPN glycoforms. **(D)** Enzymatic glycan- trimming reduces heterogeneity and results in simplified, analyzable native mass spectra of OPN glycoforms.

The first (N1H1) comprised α2-3,6,8,9 neuraminidase A, β1-4 galactosidase S, and β- *N*-acetylhexosaminidasef, which was expected to convert (extended) core-1 and core- 2 O-glycans into a core-1 (T antigen) structure, whereas extended core-3 O-glycans were expected to be processed differently. In the second combination (N1), β1-4 galactosidase S was replaced with β1-3,4 galactosidase, resulting in the further trimming of susceptible glycans to a single GalNAc residue (Tn antigen). Finally, addition of *O*-glycosidase to the N1H1 mixture (combination x) was expected to remove the remaining core-1 O-glycans, yielding a fully deglycosylated protein.

Consistent with this strategy and our core-2 hypothesis, treatment of OPN with combination N1H1 substantially simplified the native mass spectrum, yielding several sharp, well-resolved peaks (Figure 2D). These peaks could be deconvolved to yield masses of 35,540 Da, 27,616 Da and 22,136 Da that we assigned as OPN FL, OPN T1, and OPN T2, respectively. Comparison of the measured masses with the corresponding sequence-predicted masses determined by nTDMS^21^ revealed mass differences of 1,826 Da, 1,826 Da and 1,824 Da, respectively. These mass differences are in excellent agreement with the mass of five core-1 O-glycans, consisting of GalNAc (203 Da) and Gal (162 Da) residues, with a total combined mass of 1,825 Da. These results indicate that intact OPN predominantly carries a mixed population of five core-1 and core-2 O-glycans. Moreover, the fact that this glycan mass difference is conserved across OPN FL, OPN T1, and OPN T2, despite their differing C-terminal truncation points, further indicates that all five O-glycosylation sites lie N-terminal to both truncation sites.

Replacing β1-4 galactosidase S with β1-3,4 galactosidase (combination N1) enabled the cleavage of the β1-3 linked Gal from the T antigen, yielding single GalNAc residues. We measured molecular masses of 34,729 Da, 26,804 Da and 21,327 Da for OPN FL, OPN T1, and OPN T2, respectively, and the observed differences to the corresponding sequence-predicted masses were 1,015 Da, 1,015 Da and 1,015 Da, which is exactly the mass of five GalNAc (203 Da) residues. This observation further supports our conclusion that the majority of all three OPN versions carry five O-glycans. Finally, to achieve complete deglycosylation, *O*-glycosidase was added to the N1H1 mixture (combination x). While we could not resolve OPN T2, we measured masses of 33,714 Da and 25,791 Da for OPN FL and OPN T1. These are in excellent agreement with the sequence-predicted masses of 33,714 Da and 25,789 Da, confirming complete deglycosylation of the predominant OPN proteoforms. The absence of any residual mass shift indicates that the dominant proteoform (base peak) does not carry additional abundant PTMs.

### Complementary MS approaches reveal discordant phosphorylation occupancy in OPN

Beyond defining the O-glycosylation state of OPN, exoglycosidase treatment simplified the native mass spectra sufficiently to reveal lower-abundance PTMs that were previously obscured by glycan heterogeneity. In the +10-charge state (Figure 2D, insets), we observed satellite peaks separated by +80 Da, likely corresponding to phosphorylation (+79.9663 Da) but also potentially attributable to near-isobaric sulfation (+79.9568 Da), as well as lower abundance peaks corresponding to OPN proteoforms carrying an additional sixth core-1 O-glycan. The unmodified proteoform remained the dominant species (base peak), although OPN FL proteoforms carrying up to seven +80 Da modifications were resolved, with the maximum number observed depending on spectrum quality. For intact OPN FL N1H1 (Figure 3A), we observed a modification-state distribution with up to five such modifications and an average of 1.67 +80 Da modifications per OPN molecule.

**Figure 3:**
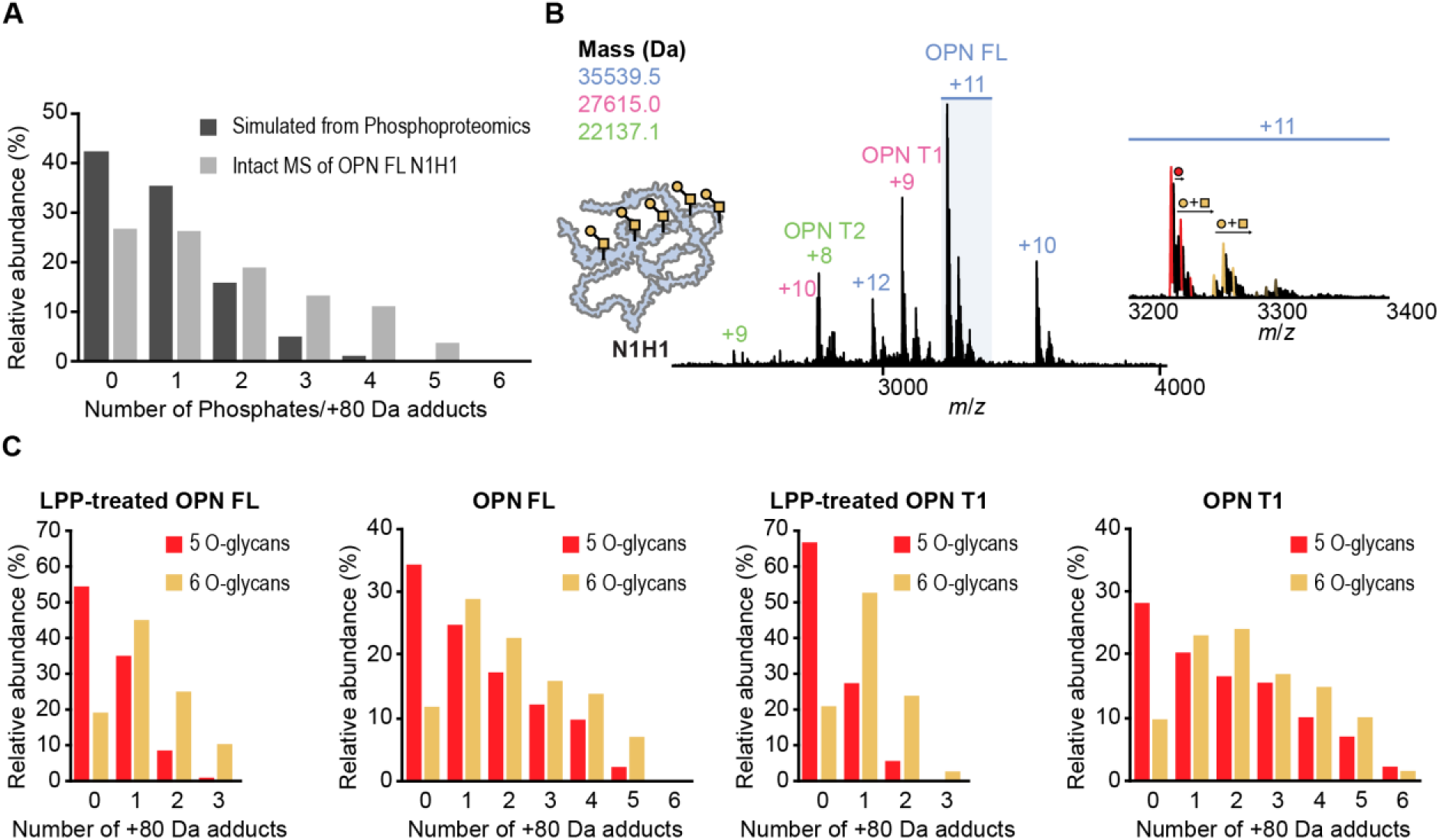
**(A)** Comparison of phosphorylation occupancy estimated from LFQ intensities by peptide-based phosphoproteomics, with the +80 Da modification-state distribution observed for intact OPN FL N1H1 by native mass spectrometry. **(B)** Native mass spectrum of glycan-trimmed and dephosphorylated OPN N1H1. **(C)** Distribution of +80 Da modifications for the predominant five-O-glycan and six-O-glycan proteoforms of intact OPN FL and OPN T1 N1H1, with and without LPP treatment.

The overall low +80 modification levels observed in HEK293-derived OPN are consistent with our previous nTDMS analysis,^21^ in which phosphorylation was detected on sequence ions from both the N- and C-termini, albeit at low frequencies. This is also comparable to phosphorylation estimates for recombinant HEK293- derived OPN in the literature ranging from none^40^ to approximately six^41^ phosphates, though substantially lower than reported levels for milk (25-26 phosphates)^22,40^ or urinary (7-8 phosphates)^40,44^ OPN.

To identify the sites underlying the +80 Da modifications observed by nMS, we performed phosphoproteomics on OPN. Our phosphoproteomics dataset covered 28 of the 41 (68%) phosphorylation sites reported in iPTMnet^56^ with a confidence score of ≥2 (Table S1). We quantified phosphorylation at 12 sites: Ser26/Ser27, Ser62, Ser63, Ser195, Ser234, Ser254, Ser263, Ser275, Ser280, Ser310, and Ser311, with the majority of phosphorylation sites located toward the C-terminus (Table S2).

To compare the phosphorylation occupancy estimated by peptide-based phosphoproteomics with that observed by nMS, minimum phosphorylation site occupancies were calculated from label-free quantification (LFQ) intensities as the ratio of the phosphorylated peptide to the total peptide signal. Summing the minimum occupancies across all quantified sites yielded an estimated average number of 0.89 phosphorylations per OPN molecule (Tables S3, S4). Comparison with the average occupancy of 1.67 calculated from nMS indicates that peptide-based phosphoproteomics does not fully quantify the +80 Da modification occupancy of OPN. Potential explanations for this discrepancy include incomplete peptide coverage, loss of labile phosphate groups during sample preparation, and analytical biases inherent to peptide-based phosphoproteomics, including the lower ionization efficiency of phosphopeptides.^57,58^ However, the +80 Da mass shifts observed by nMS are not unambiguously attributable to phosphorylation alone, raising the question of whether an additional isobaric modification also contributes to the discrepancy.^59^

### Sulfation contributes to the OPN proteoform diversity

To confirm that all +80 Da mass shifts corresponded to phosphorylation, OPN was treated with LPP and analyzed by nMS. Dephosphorylation using LPP further simplified the native mass spectra and confirmed our previous mass assignments (Figures 3B, S5). Surprisingly, we observed up to three residual +80 Da peaks for intact OPN FL N1H1 after LPP treatment (Figure 3B). For the predominant proteoforms carrying five O-glycans, the unmodified species remained the most abundant following LPP treatment, whereas among proteoforms carrying six O- glycans the singly +80 Da-modified species predominated (Figure 3C, Table S5). Importantly, this shift toward higher +80 Da occupancies in six-glycan proteoforms was not restricted to the LPP-treated samples and is therefore unlikely to be solely explained by reduced accessibility of phosphorylation sites to LPP resulting from steric hindrance by the additional O-glycan. Rather, the same trend was also observed for untreated OPN FL N1H1 as well as the truncated OPN T1 proteoforms (Figure 3C, Tables S6-S8), indicating a consistent association between the presence of an additional O-glycan and increased +80 Da modification occupancy.

The persistence of residual +80 Da species following LPP treatment, a recent meta- analysis of phosphoproteomics data highlighting OPN as a potentially sulfated protein,^60^ and a prior report of Tyr181 as a sulfation site in urinary OPN,^44^ prompted us to investigate sulfation as an additional PTM.

For the identification of sulfotyrosine-containing peptides, we performed a comparative analysis of untreated and LPP-treated OPN samples using a custom proteomics workflow^61^ that exploits the differential lability of oxygen-sulfur versus oxygen- phosphorus bonds. By applying higher-energy collisional dissociation (HCD) at low normalized collision energy (NCE) to selectively trigger loss of the labile sulfate group, we identified two candidate sulfotyrosine-containing peptides: R^176^PDIQY^181^PDATDEDITSHMESEELNGAYK^203^ and G^221^KDSY^225^ETSQLDDQSAETHSHK^241^. The latter peptide contains a single tyrosine residue, indicating Tyr225 as the sulfation site. In contrast, the former peptide contains two tyrosine residues, with Tyr181 being the more likely site of modification given its location within an acidic sequence motif characteristic of tyrosylprotein sulfotransferase (TPST)-based tyrosine sulfation.^60,61^ Our assignments were further supported by high mass accuracy, with sulfotyrosine-containing peptides showing less than 1 ppm deviation from their theoretical mass, compared to the roughly 9.5 ppm mass difference between sulfation and phosphorylation. For confident site discrimination, we additionally employed electron-transfer dissociation (ETD) and electron-transfer dissociation with supplemental HCD (EThcD) fragmentation, although −80 Da neutral loss dominated these spectra, precluding site localization; this neutral loss, however, further supports the sulfation assignment.^61^

Together, these findings support that at least a subset of the LPP-resistant +80 Da modifications detected by nMS correspond to tyrosine sulfation. Interestingly, the enrichment of residual +80 Da modifications in six-glycan proteoforms is consistent with recent reports describing cross-talk and co-localization between tyrosine sulfation and mucin-type O-glycosylation,^62^ although the mechanistic basis of this association in OPN remains to be established.

### Complementary protease strategies maximize glycosite coverage across OPN

nMS enabled us to characterize the overall glycosylation state of intact OPN but provides only partial insight into glycan macroheterogeneity (the presence or absence of glycans at specific sites) and microheterogeneity (the structural diversity of glycans at individual glycosites). Peptide-based glycoproteomics loses information about the overall glycan occupancy on the intact protein but can achieve detailed glycosite and composition analysis, and thus offers the complementary view needed to resolve glycosylation at the site level.^4,5^

To maximize protein sequence coverage and glycosite detection, we double-digested untreated OPN and exoglycosidase-treated N1H1, N1 and x using combinations of chymotrypsin (CT) and endoproteinase GluC (GluC), CT and O-glycoprotease IMPa as well as GluC and IMPa. CT and GluC were selected based on the OPN sequence to generate complementary peptide lengths suitable for LC-MS/MS analysis. IMPa is a broad O-glycan-specific endoprotease that cleaves peptide bonds immediately N-terminal to glycosylated Ser/Thr residues, accommodating a wide range of O-glycan structures, including branched and sialylated forms.^63–65^ As a result, the first residue of any IMPa-generated peptide must itself carry an O-glycan. We obtained coverage across most of the OPN sequence, including the threonine/proline-rich region that harbors a high density of previously reported glycosites (Figure 4A). Coverage was lacking for residues 87-130, which include the aspartate domain.

**Figure 4:**
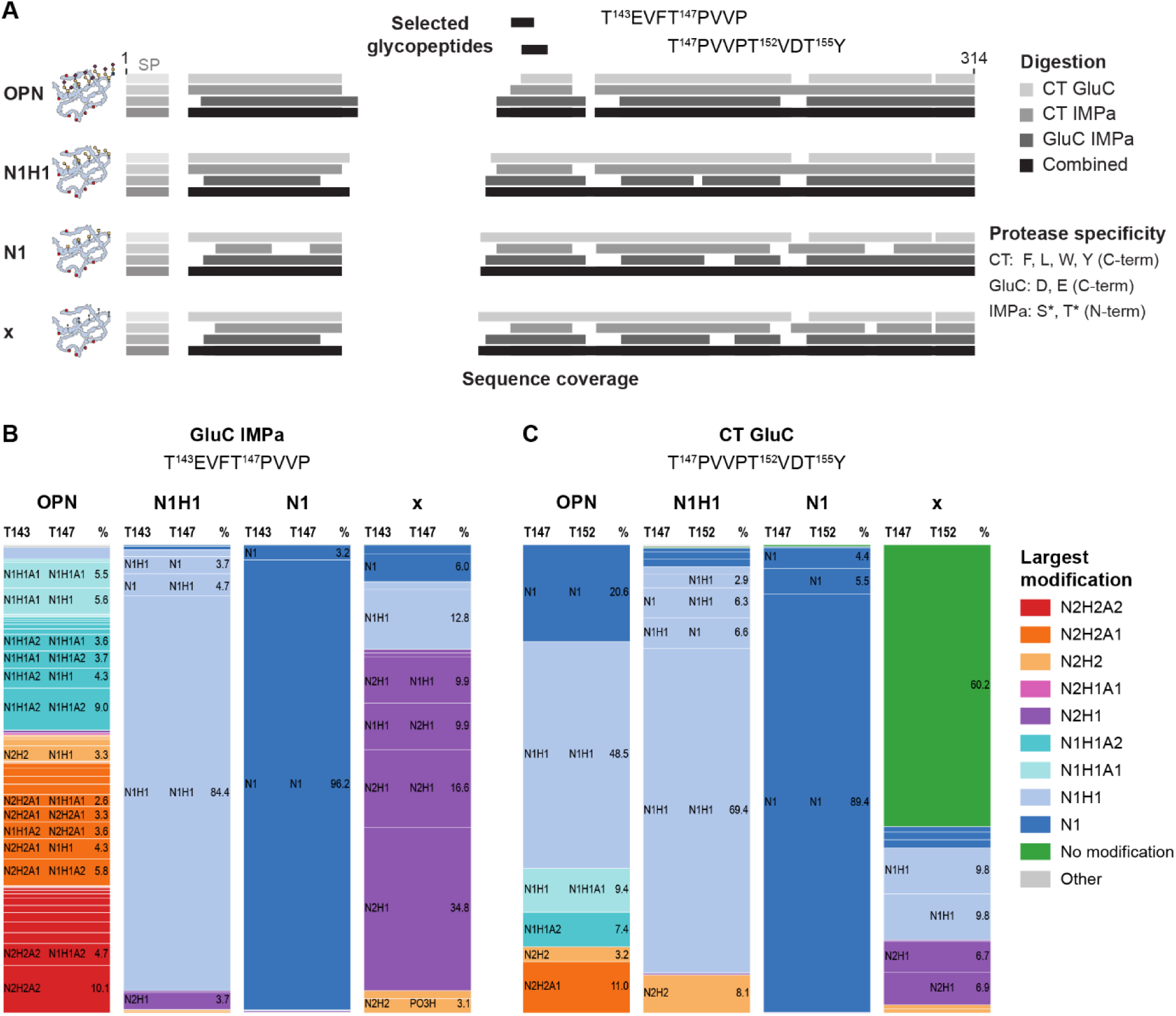
**(A)** Sequence coverage obtained in glycoproteomics with different protease combinations, with selected identified glycopeptides mapped onto the OPN sequence. **(B)** O-glycan microheterogeneity observed for Thr143 and Thr147 in the peptide TEVFTPVVP generated by GluC IMPa digestion. **(C)** O-glycan microheterogeneity observed for Thr147 and Thr152 in the peptide TPVVPTVDTY generated by CT GluC digestion. Minor fractions below 2.5% are not annotated to maintain figure clarity.

To identify and localize O-glycosylation sites, we used an optimized data-dependent HCD oxonium ion-triggered EThcD tandem MS method. In this method, detection of glycan-specific oxonium ions during HCD triggers a paired EThcD scan, required for reliable site localization since O-glycans are typically lost under collision-based dissociation alone.^8^ Data were searched in FragPipe using MSFragger-Glyco with the O-Pair module^6–8,66^, applying a 12-composition human O-glycan database against the OPN sequence. This identified several O-glycosylated peptides, predominantly within the threonine/proline-rich region (Figure S7).

### IMPa-based glycoproteomics resolves extensive glycan microheterogeneity in the threonine/proline-rich region inaccessible to conventional proteases

In both GluC IMPa and CT IMPa digests, we observed the peptide T^143^EVFT^147^PVVP (Figure 4B). Both Thr143 and Thr147 exhibit pronounced microheterogeneity, carrying predominantly sialylated core-1 as well as sialylated (extended) core-2 O-glycans. Exoglycosidase treatment (conditions N1H1 and N1) markedly reduced this glycan complexity, yielding peptides carrying either core-1 or GalNAc structures, respectively. These simplified glycoforms were identified using the same search parameters and glycan database applied to the untreated sample, suggesting that the observed reduction in complexity reflects genuine enzymatic trimming rather than a search-space artifact. The resulting glycoform assignments also match the corresponding nMS assignments, confirming that the two orthogonal methods converge on a consistent picture of OPN glycosylation – at least for Thr143 and Thr147 under simplified glycan conditions. Whether this agreement extends to the untreated, more heterogeneous sialylated glycan populations at these or other sites is a question we return to below. Unexpectedly, the corresponding peptide from the sample treated with glycosidases to achieve complete deglycosylation (condition x) still carried predominantly core-2 glycans. This apparent contradiction is explained by a selection bias inherent to IMPa digestion: because IMPa can only cleave at an O-glycosylated residue, any site where deglycosylation succeeded is no longer an IMPa substrate, and therefore yields no peptide from IMPa treatment. As a result, IMPa digestion ofthis sample necessarily enriches for the minority of sites where glycan removal was incomplete, rather than reflecting the sample’s overall (de)glycosylation state. This bias likely explains the predominance of core-2 in the observed peptide, since bulkier core-2 structures are plausibly more resistant to complete exoglycosidase trimming than core-1.

Comparison with a related peptide, T^147^PVVPT^152^VDT^155^Y, identified in the CT GluC digestion yielded additional insight (Figure 4C). In the deglycosylated condition (x), this peptide was observed primarily in its deglycosylated form, with only residual O- glycans detectable, consistent with the nMS data. The peptide from untreated OPN digested by CT GluC showed markedly less microheterogeneity, carrying shorter, less sialylated O-glycans than the related IMPa-derived peptide. This likely reflects the limited efficiency of conventional proteases at digesting extended, glycan-rich sequences, consistent with the fact that T^147^PVVPT^152^VDT^155^Y was the only glycopeptide identified from the threonine/proline-rich region by CT GluC digestion.

Thr152 was also covered by glycopeptides from GluC IMPa and CT IMPa digestion (Figures S8, S12), showing less pronounced microheterogeneity than Thr143 or Thr147 and carrying predominantly sialylated core-1 *O*-glycans. For the peptide T^152^VDT^155^YDGRGDSVVYGLRSK, our search also identified both Thr152 and Thr155 as candidate O-glycosites. However, Thr155 alone cannot account for this peptide’s generation, since IMPa-mediated cleavage requires a glycosylated N-terminal residue, implicating Thr152 as the glycosylated site responsible for peptide formation. We did not identify any glycopeptides beginning at Thr155, which could reflect either the absence of glycosylation at this site or be explained by the lack of IMPa activity when Asp occupies the P1 position.^64^

We obtained evidence for glycosylation at Thr134 and Thr138 (Figures S9-S11,S13). Glycopeptides covering these residues, however, were only detected in the exoglycosidase-treated samples, where the O-glycans were trimmed prior to protease digestion, a strategy that inevitably sacrifices information about their microheterogeneity.

The identified OPN glycosites are predominantly located within the threonine/proline- rich region at Thr134, Thr138, Thr143, Thr147, and Thr152. We also observed O- glycosylation at Thr42, Ser49, Ser76, and Thr185 at significantly lower occupancy (Figures S8, S9, S11, S16-S18), consistent with up to seven O-glycans per molecule observed by nMS. We determined the microheterogeneity of a subset of these sites, including Thr143 and Thr147, which carry mainly sialylated core-1 and sialylated extended core-2 O-glycans, and Thr152, which carries predominantly sialylated core- 1 O-glycans. Exoglycosidase treatment was required to identify Thr134 and Thr138, demonstrating that enzymatic simplification of O-glycans can substantially improve glycosite coverage in proteomics analyses. Comparison of CT GluC with IMPa- digested samples further highlighted the limited efficiency of conventional proteases in digesting extended, extensively glycosylated regions. IMPa markedly improved both glycosite coverage in this region and the capture of microheterogeneity. This may also explain why earlier LC-MS/MS-based studies^41,45^ struggled to identify glycosites and characterize their microheterogeneity within the threonine/proline-rich region. A combination of Edman degradation and MALDI-MS previously identified glycosites at Thr134, Thr138, Thr143, Thr147, and Thr152 in OPN from milk and urine, but provided no detailed information on the attached glycan structures, although disialylated core- 1 O-glycosylation was inferred for urinary OPN from MALDI-MS masses.^22,44^ LC- MS/MS-based studies of OPN from HEK293 cells^41,45^ identified various other glycosylated residues or peptides, including some covering Thr42, Ser49, Ser76, and Thr185, where we observed O-glycosylation at low occupancy. For these glycopeptides, the authors observed a heterogeneous mixture of core-1, sialylated core-1, and sialylated extended core-2 O-glycans although glycosite localization was not achieved.^45^ This underscores that glycoproteomics-based approaches are more sensitive than methods relying on MALDI-MS combined with Edman degradation, enabling detection of low-occupancy glycosites that earlier approaches missed.

### Integrating native MS and glycoproteomics reveals the sialylation state of intact OPN

To quantitatively evaluate competing glycosylation hypotheses, we developed a forward simulation framework that reconstructs representative intact OPN mass distributions from experimentally determined glycan occupancy and composition. Using this framework, we tested different glycosylation models by benchmarking their predicted mass distributions against the experimental DIA-PTCR MS/MS mass envelope of untreated OPN FL. Quantitative glycan occupancy was obtained from protein-centric nMS of intact, exoglycosidase-treated OPN, while the underlying compositional heterogeneity was resolved by bottom-up glycoproteomics, together providing the experimental inputs for the framework.

Specifically, we derived the underlying protein mass distribution, including the remaining +80 Da modifications, from the nMS spectrum of OPN FL N1H1 after accounting for the mass contribution of five core-1 O-glycans (Figure 5A, Table S9). The simplified mass spectrum of LPP-treated OPN FL N1H1 was then used to determine the relative abundances of proteoforms carrying five, six, or seven O-glycans (Table S10). Different per-glycan mass distributions representing alternative glycosylation models were constructed from the experimentally determined glycoproteomic data. The underlying protein mass distribution was then convolved with the corresponding N-fold glycan mass distributions, weighted according to the observed abundances of proteoforms carrying five, six, or seven O-glycans, and summed to reconstruct the intact OPN mass distribution for each glycosylation model.

**Figure 5:**
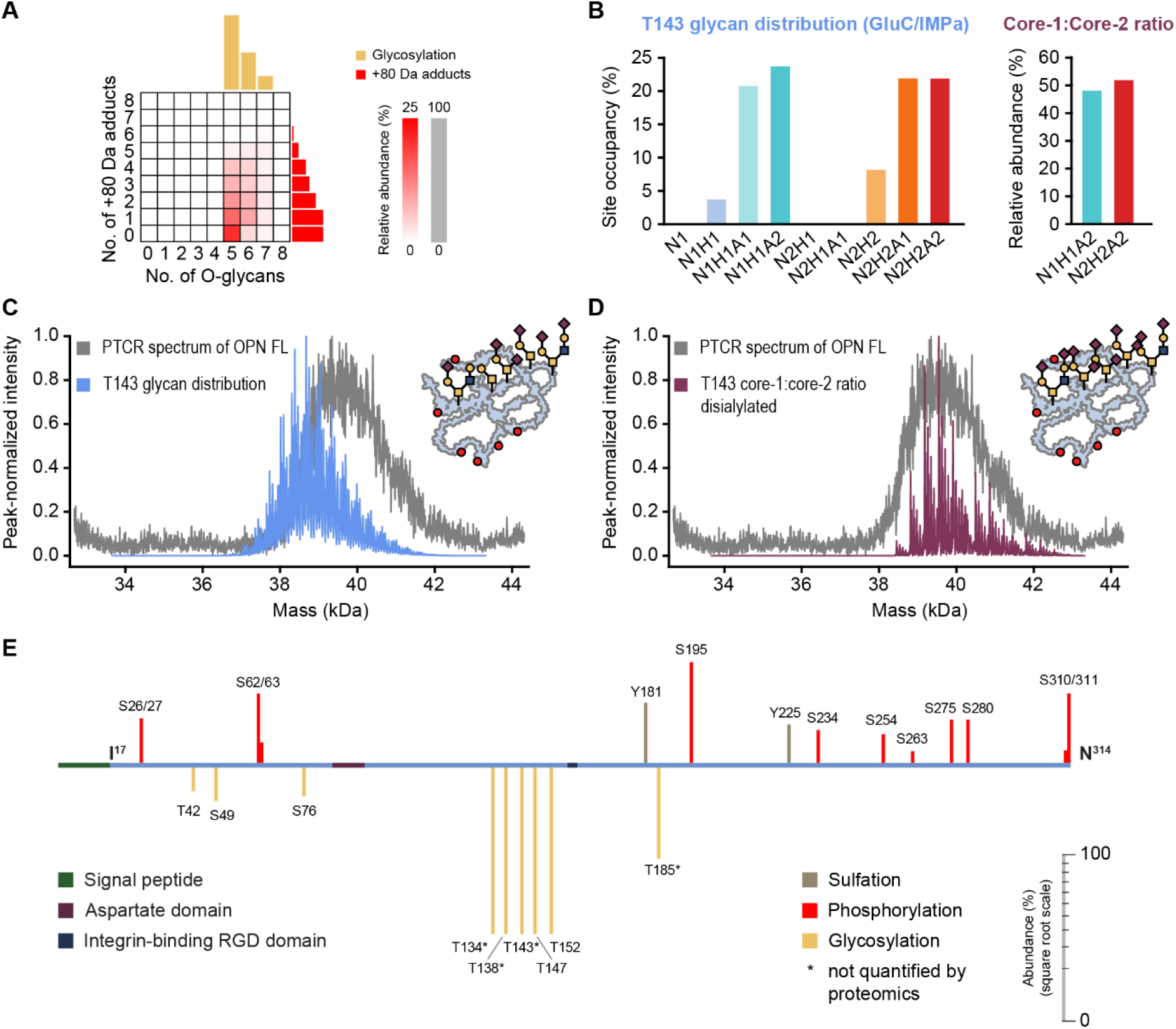
**(A)** Matrix-representation of the phosphorylation and sulfation (+80 Da adducts) as well as O-glycosylation state derived from the native mass spectrum of intact OPN FL N1H1. **(B)** Glycan composition at Thr143, determined by glycoproteomic analysis of the peptide T^143^EVFT^147^PVVP following IMPa and GluC digestion of untreated OPN. Left: site occupancy for each detected glycoform, including core-1 (N1H1, N1H1A1, N1H1A2) and core-2 (N2H1, N2H1A1, N2H2, N2H2A1, N2H2A2) compositions. Right: relative abundance assuming disialylation of core-1 (N1H1A2) and core-2 (N2H2A2) glycoforms. Comparison of simulated intact OPN mass distributions with the DIA-PTCR MS/MS mass envelope of untreated OPN FL. Simulations assume **(C)** the measured glycan composition distribution of Thr143 or **(D)** exclusively disialylated glycans with the experimental Thr143 core-1:core-2 ratio. (**E**) Post-translational modifications of OPN identified in this work.

The first model incorporated the experimentally determined per-glycan composition distribution and core-1:core-2 ratio for Thr143, derived from glycoproteomic analysis of the peptide T^143^EVFT^147^PVVP following IMPa and GluC digestion of untreated OPN (Figure 5B). We selected Thr143 for this model because it displayed the most extensive glycan profile of any site analyzed, including a high abundance of sialylated glycans, while producing consistent glycan distributions and core-1:core-2 ratios across both IMPa GluC and IMPa CT digestion workflows (Tables S11, S12).

Despite using the most extensively glycosylated site as input, the resulting simulated mass distribution still underestimated the experimental PTCR spectrum (Figure 5C). This discrepancy mirrors recent findings for N-glycosylated proteins, where bottom-up glycoproteomics has similarly been shown to underestimate intact mass distributions measured by nMS or charge-detection MS.^10,11^ In those studies, glycopeptide-derived mass distributions were shifted toward lower molecular weights relative to the corresponding intact-MS spectra, an effect attributed to an underrepresentation of highly sialylated glycans in the bottom-up data. This bias has been explained by reduced ionization efficiencies of highly sialylated glycopeptides and the lability of sialic acid residues during ionization and activation, resulting in an underestimation of larger, more highly sialylated glycan species.

To account for the potential underrepresentation of sialylation in the glycoproteomics data, we constructed a second model in which every glycan was assumed to be disialylated (N1H1A2 and N2H2A2), while keeping the experimentally determined core-1:core-2 ratio for Thr143 unchanged. Comparing this fully disialylated model against the experimental PTCR distribution therefore provides a direct test of whether incomplete capture of glycan sialylation is sufficient to explain the original discrepancy. The disialylated model substantially improved agreement between the simulated and experimental PTCR mass distributions (Figure 5D), indicating that underrepresentation of highly sialylated glycans is a major contributor to the discrepancy between bottom-up glycoproteomics and intact MS.

Comparing the Thr143 glycan distribution model with the disialylated model showed that the experimental PTCR spectrum retained a low-mass population not captured by the disialylated model, indicating that a small proportion of O-glycans are likely not fully sialylated. Together, these findings suggest that the experimentally observed OPN glycosylation is best described by an intermediate composition, comprising predominantly disialylated core-1 and disialylated extended core-2 glycans, alongside a minor proportion of shorter, less extensively sialylated glycans. To quantify this proportion, we performed additional simulations in which the per-glycan sialylation fraction was systematically increased from 0% to 100% in 10% steps (Figure S19A). The simulated distribution best matched the experimental PTCR spectrum at approximately 90% overall sialylation (Figure S19B), consistent with a minor subpopulation of glycans that are not fully disialylated. Incorporating sodium adduction into the models increased peak broadening but did not change the qualitative agreement between the simulated and experimental mass distributions, nor the conclusions drawn from the different glycosylation models (Figure S20).

Finally, these results demonstrate the complementarity of nMS and glycoproteomics for the characterization of highly O-glycosylated proteins. Conventional proteases alone fail to adequately capture the extensive glycan heterogeneity of OPN. This limitation can be overcome through the use of glycoproteases such as IMPa, which preserve substantially greater glycan compositional and structural diversity. However, because IMPa cleavage depends on the presence of O-glycans, it cannot be used to accurately determine site-specific glycosylation occupancies from proteomics alone, as non-glycosylated sites are not cleaved. By integrating the glycan composition profiles obtained from IMPa-based glycoproteomics with glycan occupancy information derived from nMS following exoglycosidase treatment, we developed a framework for estimating and evaluating plausible intact OPN glycoform distributions that could not be obtained from either approach independently.

## Conclusions

Heterogeneously O-glycosylated proteins with additional co-occurring PTMs remain among the most challenging biomolecules to characterize by current analytical methods. Here, we developed an integrated MS strategy, combining protein-centric nMS, glycoproteomics, and forward simulation, to characterize the PTM landscape of human OPN, a disease-associated, extensively post-translationally modified extracellular matrix protein.

DIA-PTCR MS/MS revealed three proteoform populations corresponding to OPN FL and two truncations, with approximate masses of 39.5 kDa, 31.6 kDa, and 26.0 kDa. Native MS and glycoproteomics demonstrated that OPN FL and its truncations predominantly carried five sialylated core-1 and sialylated extended core-2 O-glycans, while a minor OPN FL population carried up to seven O-glycans. No evidence for N-glycosylation was observed. Glycoproteomics identified Thr134, Thr138, Thr143, Thr147, and Thr152 as major glycosites, alongside Thr42, Ser49, Ser76, and Thr185, at significantly lower occupancy (Figure 5E), and resolved the microheterogeneity of Thr143, Thr147, and Thr152 as predominantly sialylated core-1 and sialylated extended core-2 O-glycans.

Integrating nMS and glycoproteomics into a forward simulation allowed us to reconstruct representative intact OPN proteoform distributions, which we validated by benchmarking against the intact-mass distribution obtained by PTCR MS. Critically, this comparison revealed that bottom-up glycoproteomics alone systematically underestimates the true extent of OPN modification, consistent with an underrepresentation of highly sialylated glycans in bottom-up glycoproteomics. Assuming complete glycan sialylation, while preserving the experimentally determined core-1:core-2 ratio, substantially improved agreement between the simulated and experimental mass distributions.

The predominant OPN proteoform lacked detectable +80 Da modifications, though proteoforms carrying up to seven such modifications were observed by nMS. Phosphoproteomics quantified 12 known phosphorylation sites, predominantly within the C-terminal region, together with two candidate sulfation sites, Tyr181 and Tyr225. Collectively, these findings indicate that human HEK293-expressed OPN is less extensively phosphorylated and O-glycosylated than previously reported, sharing several features with urinary OPN, including truncations, relatively low phosphorylation levels, and O-glycosylation predominantly within the threonine/proline-rich region.

Together, these results demonstrate the power of our integrated MS strategy. Native MS combined with sequential exoglycosidase treatment resolved OPN’s amino acid truncation, phosphorylation, sulfation, and O-glycosylation state directly on the intact protein, while glycoproteomics with tailored (glyco)protease combinations localized glycosylation sites and glycan types inaccessible to conventional trypsin-based approaches. Only by integrating information from both datasets through forward simulation and comparing the result with the intact-mass distribution obtained by PTCR MS could we uncover the systematic underestimation of glycan sialylation observed in bottom-up analysis alone. This integrated strategy provides an in-depth, quantitative view of OPN’s proteoform diversity and will facilitate future studies of how PTM alterations contribute to OPN’s role in disease. Beyond OPN, we have established a broadly applicable framework for resolving proteoform diversity in other extensively O-glycosylated proteins.

## Acknowledgements

The authors thank Di Wu for helpful discussion. K.C.Z., E.H., and J.L.P.B. were supported by the Wellcome Leap program DeltaTissue and the BBSRC (BB/W00349X/1). J.L.B., C.A.L., and C.V.R. were supported by the European Research Council (ERC) Advanced Grant (EP/Y029259/1) and by the Wellcome Leap program Multi-Channel Psych. L.A.D. and C.E.E. were supported by the BBSRC (BB/R000182/1 and BB/W00349X/1). S.A.B. was supported by a Wellcome Early-Career Award (312750/Z/24/Z), and W.B.S. by a UK Research and Innovation Future Leaders Fellowship (MR/V02213X/1).

## Author contributions

K.C.Z. and J.L.P.B. conceived and designed the study; K.C.Z. and E.H. prepared protein samples; K.C.Z. and J.L.B. performed native MS experiments and analyzed data; L.A.D. performed phosphoproteomics and sulfoproteomics sample preparation, LC-MS/MS, and data analysis; K.C.Z., J.L.B., and S.A.B. performed glycoproteomics sample preparation, LC-MS/MS, and data analysis; K.C.Z. and J.L.B. developed the simulation framework; C.A.L., C.E.E., C.V.R., W.B.S., and J.L.P.B. supervised the research and acquired funding; K.C.Z., J.L.B., W.B.S., and J.L.P.B. wrote the original manuscript draft; all authors contributed to data interpretation and reviewed, edited, and approved the final manuscript.

## References

(1) Moremen, K. W.; Tiemeyer, M.; Nairn, A. V. Vertebrate Protein Glycosylation: Diversity, Synthesis and Function. Nat. Rev. Mol. Cell Biol. 2012, 13 (7), 448–462. 10.1038/nrm3383

(2) Reily, C.; Stewart, T. J.; Renfrow, M. B.; Novak, J. Glycosylation in Health and Disease. Nat. Rev. Nephrol. 2019, 15 (6), 346–366. 10.1038/s41581-019-0129-4

(3) Wandall, H. H.; Nielsen, M. A. I.; King-Smith, S.; de Haan, N.; Bagdonaite, I. Global Functions of O-Glycosylation: Promises and Challenges in O-Glycobiology. FEBS J. 2021, 288 (24), 7183– 7212. 10.1111/febs.16148

(4) Bagdonaite, I.; Malaker, S. A.; Polasky, D. A.; Riley, N. M.; Schjoldager, K.; Vakhrushev, S. Y.; Halim, A.; Aoki-Kinoshita, K. F.; Nesvizhskii, A. I.; Bertozzi, C. R.; Wandall, H. H.; Parker, B. L.; Thaysen-Andersen, M.; Scott, N. E., Glycoproteomics. Nat. Rev. Methods Primer 2022, 2 (1), 48. 10.1038/s43586-022-00128-4

(5) Chen, S.; Wu, D.; Robinson, C. V.; Struwe, W. B. Native Mass Spectrometry Meets Glycomics: Resolving Structural Detail and Occupancy of Glycans on Intact Glycoproteins. Anal. Chem. 2021, 93 (30), 10435–10443. 10.1021/acs.analchem.1c01460

(6) Polasky, D. A.; Yu, F.; Teo, G. C.; Nesvizhskii, A. I. Fast and Comprehensive N- and O- Glycoproteomics Analysis with MSFragger-Glyco. Nat. Methods 2020, 17 (11), 1125–1132. 10.1038/s41592-020-0967-9

(7) Lu, L.; Riley, N. M.; Shortreed, M. R.; Bertozzi, C. R.; Smith, L. M. O-Pair Search with MetaMorpheus for O-Glycopeptide Characterization. Nat. Methods 2020, 17 (11), 1133–1138. 10.1038/s41592-020-00985-5

(8) Riley, N. M.; Bertozzi, C. R. Deciphering O-Glycoprotease Substrate Preferences with O-Pair Search. *Mol*. Omics 2022, 18 (10), 908–922. 10.1039/D2MO00244B

(9) Yang, Y.; Liu, F.; Franc, V.; Halim, L. A.; Schellekens, H.; Heck, A. J. R. Hybrid Mass Spectrometry Approaches in Glycoprotein Analysis and Their Usage in Scoring Biosimilarity. Nat. Commun. 2016, 7 (1), 13397. 10.1038/ncomms13397

(10) Čaval, T.; Buettner, A.; Haberger, M.; Reusch, D.; Heck, A. J. R. Discrepancies between High- Resolution Native and Glycopeptide-Centric Mass Spectrometric Approaches: A Case Study into the Glycosylation of Erythropoietin Variants. J. Am. Soc. Mass Spectrom. 2021, 32 (8), 2099– 2104. 10.1021/jasms.1c00060

(11) Miller, L. M.; Barnes, L. F.; Raab, S. A.; Draper, B. E.; El-Baba, T. J.; Lutomski, C. A.; Robinson, C. V.; Clemmer, D. E.; Jarrold, M. F. Heterogeneity of Glycan Processing on Trimeric SARS-CoV-2 Spike Protein Revealed by Charge Detection Mass Spectrometry. J. Am. Chem. Soc. 2021, 143 (10), 3959–3966. 10.1021/jacs.1c00353

(12) Wu, D.; Struwe, W. B.; Harvey, D. J.; Ferguson, M. A. J.; Robinson, C. V. N-Glycan Microheterogeneity Regulates Interactions of Plasma Proteins. Proc. Natl. Acad. Sci. 2018, 115 (35), 8763–8768. 10.1073/pnas.1807439115

(13) Po, A.; Eyers, C. E. Top-Down Proteomics and the Challenges of True Proteoform Characterization. J. Proteome Res. 2023, 22 (12), 3663–3675. 10.1021/acs.jproteome.3c00416

(14) Rosati, S.; Rose, R. J.; Thompson, N. J.; van Duijn, E.; Damoc, E.; Denisov, E.; Makarov, A.; Heck, A. J. R. Exploring an Orbitrap Analyzer for the Characterization of Intact Antibodies by Native Mass Spectrometry. Angew. Chem. Int. Ed. 2012, 51 (52), 12992–12996. 10.1002/anie.201206745

(15) Yang, Y.; Barendregt, A.; Kamerling, J. P.; Heck, A. J. R. Analyzing Protein Micro-Heterogeneity in Chicken Ovalbumin by High-Resolution Native Mass Spectrometry Exposes Qualitatively and Semi-Quantitatively 59 Proteoforms. Anal. Chem. 2013, 85 (24), 12037–12045. 10.1021/ac403057y

(16) Yen, H.-Y.; Liko, I.; Gault, J.; Wu, D.; Struwe, W. B.; Robinson, C. V. Correlating Glycoforms of DC-SIGN with Stability Using a Combination of Enzymatic Digestion and Ion Mobility Mass Spectrometry. Angew. Chem. 2020, 132 (36), 15690–15694. 10.1002/ange.202005727

(17) Rolland, A. D.; Prell, J. S. Approaches to Heterogeneity in Native Mass Spectrometry. Chem. Rev. 2022, 122 (8), 7909–7951. 10.1021/acs.chemrev.1c00696

(18) Schachner, L. F.; Mullen, C.; Phung, W.; Hinkle, J. D.; Beardsley, M. I.; Bentley, T.; Day, P.; Tsai, C.; Sukumaran, S.; Baginski, T.; DiCara, D.; Agard, N. J.; Masureel, M.; Gober, J.; ElSohly, A. M.; Melani, R.; Syka, J. E. P.; Huguet, R.; Marty, M. T.; Sandoval, W. Exposing the Molecular Heterogeneity of Glycosylated Biotherapeutics. Nat. Commun. 2024, 15 (1), 3259. 10.1038/s41467-024-47693-8

(19) Wohlschlager, T.; Scheffler, K.; Forstenlehner, I. C.; Skala, W.; Senn, S.; Damoc, E.; Holzmann, J.; Huber, C. G. Native Mass Spectrometry Combined with Enzymatic Dissection Unravels Glycoform Heterogeneity of Biopharmaceuticals. Nat. Commun. 2018, 9 (1), 1713. 10.1038/s41467-018-04061-7

(20) Roberts, D. S.; Mann, M.; Melby, J. A.; Larson, E. J.; Zhu, Y.; Brasier, A. R.; Jin, S.; Ge, Y. Structural O-Glycoform Heterogeneity of the SARS-CoV-2 Spike Protein Receptor-Binding Domain Revealed by Top-Down Mass Spectrometry. J. Am. Chem. Soc. 2021, 143 (31), 12014– 12024. 10.1021/jacs.1c02713

(21) Bennett, J. L.; El-Baba, T. J.; Zouboulis, K. C.; Kirschbaum, C.; Song, H.; Butroid, F. I.; Benesch, J. L. P.; Lutomski, C. A.; Robinson, C. V. Uncovering Hidden Protein Modifications with Native Top-down Mass Spectrometry. Nat. Methods 2025, 22 (10), 2127–2137. 10.1038/s41592-025-02846-5

(22) Christensen, B.; Nielsen, M. S.; Haselmann, K. F.; Petersen, T. E.; Sørensen, E. S. Post-Translationally Modified Residues of Native Human Osteopontin Are Located in Clusters: Identification of 36 Phosphorylation and Five O-Glycosylation Sites and Their Biological Implications. Biochem. J. 2005, 390 (1), 285–292. 10.1042/BJ20050341

(23) Bellahcène, A.; Castronovo, V.; Ogbureke, K. U. E.; Fisher, L. W.; Fedarko, N. S. Small Integrin-Binding Ligand N-Linked Glycoproteins (SIBLINGs): Multifunctional Proteins in Cancer. Nat. Rev. Cancer 2008, 8 (3), 212–226. 10.1038/nrc2345

(24) Platzer, G.; Schedlbauer, A.; Chemelli, A.; Ozdowy, P.; Coudevylle, N.; Auer, R.; Kontaxis, G.; Hartl, M.; Miles, A. J.; Wallace, B. A.; Glatter, O.; Bister, K.; Konrat, R. The Metastasis-Associated Extracellular Matrix Protein Osteopontin Forms Transient Structure in Ligand Interaction Sites. Biochemistry 2011, 50 (27), 6113–6124. 10.1021/bi200291e

(25) Kurzbach, D.; Platzer, G.; Schwarz, T. C.; Henen, M. A.; Konrat, R.; Hinderberger, D. Cooperative Unfolding of Compact Conformations of the Intrinsically Disordered Protein Osteopontin. Biochemistry 2013, 52 (31), 5167–5175. 10.1021/bi400502c

(26) Weber, G. F.; Ashkar, S.; Glimcher, M. J.; Cantor, H. Receptor-Ligand Interaction Between CD44 and Osteopontin (Eta-1). Science 1996, 271 (5248), 509–512. 10.1126/science.271.5248.509

(27) Ashkar, S.; Weber, G. F.; Panoutsakopoulou, V.; Sanchirico, M. E.; Jansson, M.; Zawaideh, S.; Rittling, S. R.; Denhardt, D. T.; Glimcher, M. J.; Cantor, H. Eta-1 (Osteopontin): An Early Component of Type-1 (Cell-Mediated) Immunity. Science 2000, 287 (5454), 860–864. 10.1126/science.287.5454.860

(28) Chellaiah, M. A.; Biswas, R. S.; Rittling, S. R.; Denhardt, D. T.; Hruska, K. A. Rho-Dependent Rho Kinase Activation Increases CD44 Surface Expression and Bone Resorption in Osteoclasts*. J. Biol. Chem. 2003, 278 (31), 29086–29097. 10.1074/jbc.M211074200

(29) Moorman, H. R.; Poschel, D.; Klement, J. D.; Lu, C.; Redd, P. S.; Liu, K. Osteopontin: A Key Regulator of Tumor Progression and Immunomodulation. Cancers 2020, 12 (11), 3379. 10.3390/cancers12113379

(30) Lin, E. Y.-H.; Xi, W.; Aggarwal, N.; Shinohara, M. L. Osteopontin (OPN)/SPP1: From Its Biochemistry to Biological Functions in the Innate Immune System and the Central Nervous System (CNS). Int. Immunol. 2023, 35 (4), 171–180. 10.1093/intimm/dxac060

(31) Panda, V. K.; Mishra, B.; Nath, A. N.; Butti, R.; Yadav, A. S.; Malhotra, D.; Khanra, S.; Mahapatra, S.; Mishra, P.; Swain, B.; Majhi, S.; Kumari, K.; Radharani, N. N. V.; Kundu, G. C. Osteopontin: A Key Multifaceted Regulator in Tumor Progression and Immunomodulation. Biomedicines 2024, 12 (7), 1527. 10.3390/biomedicines12071527

(32) Cohen, J. D.; Li, L.; Wang, Y.; Thoburn, C.; Afsari, B.; Danilova, L.; Douville, C.; Javed, A. A.; Wong, F.; Mattox, A.; Hruban, R. H.; Wolfgang, C. L.; Goggins, M. G.; Dal Molin, M.; Wang, T.-L.; Roden, R.; Klein, A. P.; Ptak, J.; Dobbyn, L.; Schaefer, J.; Silliman, N.; Popoli, M.; Vogelstein, J. T.; Browne, J. D.; Schoen, R. E.; Brand, R. E.; Tie, J.; Gibbs, P.; Wong, H.-L.; Mansfield, A. S.; Jen, J.; Hanash, S. M.; Falconi, M.; Allen, P. J.; Zhou, S.; Bettegowda, C.; Diaz, L. A.; Tomasetti, C.; Kinzler, K. W.; Vogelstein, B.; Lennon, A. M.; Papadopoulos, N. Detection and Localization of Surgically Resectable Cancers with a Multi-Analyte Blood Test. Science 2018, 359 (6378), 926– 930. 10.1126/science.aar3247

(33) Gu, Y.; Taifour, T.; Bui, T.; Zuo, D.; Pacis, A.; Poirier, A.; Attalla, S.; Fortier, A.-M.; Sanguin-Gendreau, V.; Pan, T.-C.; Papavasiliou, V.; Lin, N. U.; Hughes, M. E.; Smith, K.; Park, M.; Tremblay, M. L.; Chodosh, L. A.; Jeselsohn, R.; Muller, W. J. Osteopontin Is a Therapeutic Target That Drives Breast Cancer Recurrence. Nat. Commun. 2024, 15 (1), 9174. 10.1038/s41467-024-53023-9

(34) Gao, W.; Liu, D.; Sun, H.; Shao, Z.; Shi, P.; Li, T.; Yin, S.; Zhu, T. SPP1 Is a Prognostic Related Biomarker and Correlated with Tumor-Infiltrating Immune Cells in Ovarian Cancer. BMC Cancer 2022, 22 (1), 1367. 10.1186/s12885-022-10485-8

(35) Kariya, Y.; Kariya, Y. Osteopontin in Cancer: Mechanisms and Therapeutic Targets. *Int*. J. Transl. Med. 2022, 2 (3), 419–447. 10.3390/ijtm2030033

(36) Leung, L. L.; Myles, T.; Morser, J. Thrombin Cleavage of Osteopontin and the Host Anti-Tumor Immune Response. Cancers 2023, 15 (13), 3480. 10.3390/cancers15133480

(37) Li, H.; Lan, L.; Chen, H.; Zaw Thin, M.; Ps, H.; Nelson, J. K.; Evans, I. M.; Ruiz, E. J.; Cheng, R.; Tran, L.; Allen, M.; Ma, J.; Yi, T.; Wang, C.; He, Y.; Guppy, N.; Sadanandam, A.; Lin, S.-Z.; Zhang, C.; Behrens, A. SPP1 Is Required for Maintaining Mesenchymal Cell Fate in Pancreatic Cancer. Nature 2025, 648 (8092), 203–209. 10.1038/s41586-025-09574-y

(38) Lenos, K. J.; Miedema, D. M.; Lodestijn, S. C.; Nijman, L. E.; van den Bosch, T.; Romero Ros, X.; Lourenço, F. C.; Lecca, M. C.; van der Heijden, M.; van Neerven, S. M.; van Oort, A.; Leveille, N.; Adam, R. S.; de Sousa E Melo, F.; Otten, J.; Veerman, P.; Hypolite, G.; Koens, L.; Lyons, S. K.; Stassi, G.; Winton, D. J.; Medema, J. P.; Morrissey, E.; Bijlsma, M. F.; Vermeulen, L. Stem Cell Functionality Is Microenvironmentally Defined during Tumour Expansion and Therapy Response in Colon Cancer. Nat. Cell Biol. 2018, 20 (10), 1193–1202. 10.1038/s41556-018-0179-z

(39) Shanmugam, V.; Chackalaparampil, I.; Kundu, G. C.; Mukherjee, A. B.; Mukherjee, B. B. Altered Sialylation of Osteopontin Prevents Its Receptor-Mediated Binding on the Surface of Oncogenically Transformed tsB77 Cells. Biochemistry 1997, 36 (19), 5729–5738. 10.1021/bi961687w

(40) Christensen, B.; Kläning, E.; Nielsen, M. S.; Andersen, M. H.; Sørensen, E. S. C-Terminal Modification of Osteopontin Inhibits Interaction with the αVβ3-Integrin*. J. Biol. Chem. 2012, 287 (6), 3788–3797. 10.1074/jbc.M111.277996

(41) Kariya, Y.; Kanno, M.; Matsumoto-Morita, K.; Konno, M.; Yamaguchi, Y.; Hashimoto, Y. Osteopontin O-Glycosylation Contributes to Its Phosphorylation and Cell-Adhesion Properties. Biochem. J. 2014, 463 (1), 93–102. 10.1042/BJ20140060

(42) Gu, Y.; Muller, W. J. The Multifaceted Role of Osteopontin in Modulating the Tumor Microenvironment. Cancer Res. 2025, 85 (21), 4049–4061. 10.1158/0008-5472.CAN-25-1486

(43) Masuda, K.; Takahashi, N.; Tsukamoto, Y.; Honma, H.; Kohri, K. *N*-Glycan Structures of an Osteopontin from Human Bone. Biochem. Biophys. Res. Commun. 2000, 268 (3), 814–817. 10.1006/bbrc.2000.2224

(44) Christensen, B.; Petersen, T. E.; Sørensen, E. S. Post-Translational Modification and Proteolytic Processing of Urinary Osteopontin. Biochem. J. 2008, 411 (1), 53–61. 10.1042/BJ20071021

(45) Li, H.; Shen, H.; Yan, G.; Zhang, Y.; Liu, M.; Fang, P.; Yu, H.; Yang, P. Site-Specific Structural Characterization of O-Glycosylation and Identification of Phosphorylation Sites of Recombinant Osteopontin. Biochim. Biophys. Acta BBA - Proteins Proteomics 2015, 1854 (6), 581–591. 10.1016/j.bbapap.2014.09.025

(46) Oyama, M.; Kariya, Y.; Kariya, Y.; Matsumoto, K.; Kanno, M.; Yamaguchi, Y.; Hashimoto, Y. Biological Role of Site-Specific O-Glycosylation in Cell Adhesion Activity and Phosphorylation of Osteopontin. Biochem. J. 2018, 475 (9), 1583–1595. 10.1042/BCJ20170205

(47) Miura, Y.; Kato, K.; Takegawa, Y.; Kurogochi, M.; Furukawa, J.; Shinohara, Y.; Nagahori, N.; Amano, M.; Hinou, H.; Nishimura, S.-I. Glycoblotting-Assisted O-Glycomics: Ammonium Carbamate Allows for Highly Efficient O-Glycan Release from Glycoproteins. Anal. Chem. 2010, 82 (24), 10021–10029. 10.1021/ac101599p

(48) Froehlich, J. W.; Chu, C. S.; Tang, N.; Waddell, K.; Grimm, R.; Lebrilla, C. B. Label-Free Liquid Chromatography–Tandem Mass Spectrometry Analysis with Automated Phosphopeptide Enrichment Reveals Dynamic Human Milk Protein Phosphorylation during Lactation. Anal. Biochem. 2011, 408 (1), 136–146. 10.1016/j.ab.2010.08.031

(49) Tagliabracci, V. S.; Engel, J. L.; Wen, J.; Wiley, S. E.; Worby, C. A.; Kinch, L. N.; Xiao, J.; Grishin, N. V.; Dixon, J. E. Secreted Kinase Phosphorylates Extracellular Proteins That Regulate Biomineralization. Science 2012, 336 (6085), 1150–1153. 10.1126/science.1217817

(50) Tagliabracci, V. S.; Wiley, S. E.; Guo, X.; Kinch, L. N.; Durrant, E.; Wen, J.; Xiao, J.; Cui, J.; Nguyen, K. B.; Engel, J. L.; Coon, J. J.; Grishin, N.; Pinna, L. A.; Pagliarini, D. J.; Dixon, J. E. A Single Kinase Generates the Majority of the Secreted Phosphoproteome. Cell 2015, 161 (7), 1619–1632. 10.1016/j.cell.2015.05.028

(51) Mateos, B.; Holzinger, J.; Conrad-Billroth, C.; Platzer, G.; Żerko, S.; Sealey-Cardona, M.; Anrather, D.; Koźmiński, W.; Konrat, R. Hyperphosphorylation of Human Osteopontin and Its Impact on Structural Dynamics and Molecular Recognition. Biochemistry 2021, 60 (17), 1347– 1355. 10.1021/acs.biochem.1c00050

(52) Olson, F. J.; Bäckström, M.; Karlsson, H.; Burchell, J.; Hansson, G. C. A MUC1 Tandem Repeat Reporter Protein Produced in CHO-K1 Cells Has Sialylated Core 1 O-Glycans and Becomes More Densely Glycosylated If Coexpressed with Polypeptide-GalNAc-T4 Transferase. Glycobiology 2005, 15 (2), 177–191. 10.1093/glycob/cwh158

(53) Jaroentomeechai, T.; Karlsson, R.; Goerdeler, F.; Teoh, F. K. Y.; Grønset, M. N.; de Wit, D.; Chen, Y.-H.; Furukawa, S.; Psomiadou, V.; Hurtado-Guerrero, R.; Vidal-Calvo, E. E.; Salanti, A.; Boltje, T. J.; van den Bos, L. J.; Wunder, C.; Johannes, L.; Schjoldager, K. T.; Joshi, H. J.; Miller, R. L.; Clausen, H.; Vakhrushev, S. Y.; Narimatsu, Y. Mammalian Cell-Based Production of Glycans, Glycopeptides and Glycomodules. Nat. Commun. 2024, 15 (1), 9668. 10.1038/s41467-024-53738-9

(54) Nason, R.; Büll, C.; Konstantinidi, A.; Sun, L.; Ye, Z.; Halim, A.; Du, W.; Sørensen, D. M.; Durbesson, F.; Furukawa, S.; Mandel, U.; Joshi, H. J.; Dworkin, L. A.; Hansen, L.; David, L.; Iverson, T. M.; Bensing, B. A.; Sullam, P. M.; Varki, A.; de Vries, E.; de Haan, C. A. M.; Vincentelli, R.; Henrissat, B.; Vakhrushev, S. Y.; Clausen, H.; Narimatsu, Y. Display of the Human Mucinome with Defined O-Glycans by Gene Engineered Cells. Nat. Commun. 2021, 12 (1), 4070. 10.1038/s41467-021-24366-4

(55) Narimatsu, Y.; Joshi, H. J.; Nason, R.; van Coillie, J.; Karlsson, R.; Sun, L.; Ye, Z.; Chen, Y.-H.; Schjoldager, K. T.; Steentoft, C.; Furukawa, S.; Bensing, B. A.; Sullam, P. M.; Thompson, A. J.; Paulson, J. C.; Büll, C.; Adema, G. J.; Mandel, U.; Hansen, L.; Bennett, E. P.; Varki, A.; Vakhrushev, S. Y.; Yang, Z.; Clausen, H. An Atlas of Human Glycosylation Pathways Enables Display of the Human Glycome by Gene Engineered Cells. Mol. Cell 2019, 75 (2), 394–407.e5. 10.1016/j.molcel.2019.05.017

(56) Huang, H.; Arighi, C. N.; Ross, K. E.; Ren, J.; Li, G.; Chen, S.-C.; Wang, Q.; Cowart, J.; Vijay-Shanker, K.; Wu, C. H. iPTMnet: An Integrated Resource for Protein Post-Translational Modification Network Discovery. Nucleic Acids Res. 2018, 46 (D1), D542–D550. 10.1093/nar/gkx1104

(57) Janek, K.; Wenschuh, H.; Bienert, M.; Krause, E. Phosphopeptide Analysis by Positive and Negative Ion Matrix-Assisted Laser Desorption/Ionization Mass Spectrometry. Rapid Commun. Mass Spectrom. RCM 2001, 15 (17), 1593–1599. 10.1002/rcm.417

(58) Bubis, J. A.; Gorshkov, V.; Gorshkov, M. V.; Kjeldsen, F. PhosphoShield: Improving Trypsin Digestion of Phosphoproteins by Shielding the Negatively Charged Phosphate Moiety. J. Am. Soc. Mass Spectrom. 2020, 31 (10), 2053–2060. 10.1021/jasms.0c00171

(59) Daly, L. A.; Clarke, C. J.; Po, A.; Oswald, S. O.; Eyers, C. E. Considerations for Defining +80 Da Mass Shifts in Mass Spectrometry-Based Proteomics: Phosphorylation and Beyond. Chem. Commun. 2023, 59 (77), 11484–11499. 10.1039/d3cc02909c

(60) Tzvetkov, J.; Eyers, C. E.; Eyers, P. A.; Ramsbottom, K. A.; Oswald, S. O.; Harris, J. A.; Sun, Z.; Deutsch, E. W.; Jones, A. R. Searching for Sulfotyrosines (sY) in a HA(pY)STACK. J. Proteome Res. 2025, 24 (3), 1250–1264. 10.1021/acs.jproteome.4c00907

(61) Daly, L. A.; Byrne, D. P.; Perkins, S.; Brownridge, P. J.; McDonnell, E.; Jones, A. R.; Eyers, P. A.; Eyers, C. E. Custom Workflow for the Confident Identification of Sulfotyrosine-Containing Peptides and Their Discrimination from Phosphopeptides. J. Proteome Res. 2023, 22 (12), 3754–3772. 10.1021/acs.jproteome.3c00425

(62) Mehta, A. Y.; Heimburg-Molinaro, J.; Cummings, R. D.; Goth, C. K. Emerging Patterns of Tyrosine Sulfation and *O*-Glycosylation Cross-Talk and Co-Localization. Curr. Opin. Struct. Biol. 2020, 62, 102–111. 10.1016/j.sbi.2019.12.002

(63) Bardoel, B. W.; Hartsink, D.; Vughs, M. M.; de Haas, C. J. C.; van Strijp, J. A. G.; van Kessel, K. P. M. Identification of an Immunomodulating Metalloprotease of Pseudomonas Aeruginosa (IMPa). Cell. Microbiol. 2012, 14 (6), 902–913. 10.1111/j.1462-5822.2012.01765.x

(64) Vainauskas, S.; Guntz, H.; McLeod, E.; McClung, C.; Ruse, C.; Shi, X.; Taron, C. H. A Broad-Specificity O-Glycoprotease That Enables Improved Analysis of Glycoproteins and Glycopeptides Containing Intact Complex O-Glycans. Anal. Chem. 2022, 94 (2), 1060–1069. 10.1021/acs.analchem.1c04055

(65) Suttapitugsakul, S.; Matsumoto, Y.; Aryal, R. P.; Cummings, R. D. Large-Scale and Site-Specific Mapping of the Murine Brain O-Glycoproteome with IMPa. Anal. Chem. 2023, 95 (36), 13423– 13430. 10.1021/acs.analchem.3c00408

(66) Polasky, D. A.; Lu, L.; Yu, F.; Li, K.; Shortreed, M. R.; Smith, L. M.; Nesvizhskii, A. I. Quantitative Proteome-Wide O-Glycoproteomics Analysis with FragPipe. Anal. Bioanal. Chem. 2025, 417 (5), 921–930. 10.1007/s00216-024-05382-x

