## Supporting Information for "An integrated mass spectrometry strategy for quantifying the proteoform diversity of the extensively modified O-glycoprotein Osteopontin"

### Materials and Methods

#### Materials

Recombinant human osteopontin (OPNa, UniProt P10451-1) from human HEK293 cells was obtained from PeproTech (Thermo Fisher Scientific, #120-35) as a lyophilized powder. Its activity was tested in a cell adhesion assay using murine B16-F1 cells by the manufacturer, where it was shown to enhance adhesion, consistent with the function of endogenous OPN. The lyophilized OPN was redissolved in sterile water to 1 mg/mL, flash-frozen in liquid nitrogen and stored at -80 °C.

#### Enzymatic treatments

Desialylation: 9 µg OPN was incubated in 10 µL 1X GlycoBuffer 1 (50 mM sodium acetate, 5 mM $CaCl_2$, pH 5.5; New England Biolabs, NEB) with 60 units of *α*2-3,6,8,9 neuraminidase A (NEB, P0722S) in 3 µL storage buffer for 16 h at 37 °C.

Deglycosylation mix II treatment: Deglycosylation mix II (NEB, P6044S) contains a mixture of PNGase F, *O*-glycosidase, *α*2-3,6,8,9 neuraminidase A, *β*1-4 galactosidase S and *β*-*N*-acetylhexosaminidase$_f$, enabling the removal of N-glycans and many O-glycans from glycoproteins. 25 µg OPN in 40 µL water was incubated with 5 µL deglycosylation mix II and 5 µL 10X deglycosylation mix buffer 1 (NEB, B6044) for 30 min at 25 °C and then for 16 h at 37 °C.

PNGase F treatment: 18 µg OPN in 20 µL 1X GlycoBuffer 2 (50 mM sodium phosphate, pH 7.5; NEB) was incubated with 1,500 units of PNGase F (NEB, P0709S) in 3 µL storage buffer for 16 h at 37 °C.

N1H1: 25 µg OPN in 100 µL 1X GlycoBuffer 1 was incubated with 80 units of *α*2-3,6,8,9 neuraminidase A, 32 units of *β*1-4 galactosidase S (NEB, P0745S) and 20 units of *β*-*N*-acetylhexosaminidase$_f$ (NEB, P0721S) in 12 µL storage buffer for 48 h at 37 °C.

N1: 25 µg OPN in 100 µL 1X GlycoBuffer 1 was treated with 80 units of *α*2-3,6,8,9 neuraminidase A, 32 units of *β*1-3,4 galactosidase (NEB, P0746S) and 20 units of *β*-*N*-acetylhexosaminidase$_f$ in 12 µL storage buffer for 48 h at 37 °C.

x: 25 µg OPN in 100 µL 1X GlycoBuffer 2 was treated with 80 units of *α*2-3,6,8,9 neuraminidase A, 32 units of *β*1-4 galactosidase S, 20 units of *β*-*N*-acetylhexosaminidase$_f$ and 200 units of *O*-glycosidase (NEB, P0733S) in 16 µL storage buffer for 48 h at 37 °C.

Dephosphorylation: 25 µg OPN in 50 µL PMP buffer (50 mM HEPES, 100 mM NaCl, 2 mM DTT, 0.01% Brij 35, pH 7.5; NEB) supplemented with 1 mM $MnCl_2$ was incubated with 400 units of lambda protein phosphatase (LPP; NEB, P0753S) for 1.5 h to 16 h at 30 °C. To reduce the observed LPP signal intensity in native mass spectra, the amount of enzyme added prior to MS analysis was decreased ten-fold.

#### SDS-PAGE

OPN samples were analyzed by SDS-PAGE to assess glycan cleavage and dephosphorylation. 9 µL protein samples were mixed with 4X NuPAGE LDS sample buffer (Thermo Fisher Scientific) at a 3:1 sample-to-buffer ratio and resolved on precast 4%-12% NuPAGE Bis-Tris mini protein gels (Thermo Fisher Scientific) in MES SDS running buffer (Thermo Fisher Scientific) at 200 V for 40 min. PageRuler prestained protein ladder, 10 kDa - 180 kDa (Thermo Fisher Scientific) was used as a molecular weight standard. Gels were stained with InstantBlue Coomassie protein stain (Abcam) and destained in water before imaging.

#### Native mass spectrometry

##### Sample preparation

OPN samples were buffer exchanged into 1 M ammonium acetate, pH 7.0 using Amicon Ultra centrifugal filters (10 kDa MWCO; Merck) following the manufacturer's instructions.

##### Native MS

Native mass spectrometry (nMS) measurements were performed on a hybrid quadrupole-Orbitrap mass spectrometer optimized for the transmission of large-*m*/*z* analytes (Q Exactive UHMR; Thermo Fisher Scientific). Native protein ions were generated from gold-coated nanoelectrospray ionization (nESI) capillaries using the following parameters: spray voltages of 0.9 kV – 1.1 kV, capillary temperatures of 100 °C – 200 °C, in-source activation of -10 V – -100 V. Mass spectra were acquired in the Orbitrap at a resolution of 17,500 FWHM at *m*/*z* 200. The obtained mass spectra were averaged, subjected to minimal Gaussian smoothing and analyzed manually or, where applicable, using UniDec software version 5.0.1[1].

##### Proton-transfer charge-reduction MS

DIA-PTCR MS measurements were performed on an Orbitrap Ascend Structural Biology Edition Tribrid mass spectrometer equipped with a front-end glow discharge perfluoroperhydrophenanthrene (PFPP) ion source. The mass spectrometer was operated in intact protein mode and the ionization parameters were as described in the nMS section. For PTCR $MS^2$ spectra, narrow isolation windows (*m*/*z* 15) were sequentially selected via the quadrupole over the *m*/*z* range of 2,845 – 4,415. Isolated ions were reacted with PFPP for 50 ms before being transferred to the Orbitrap for mass analysis. Three 32 ms transients (resolution of 15,000 FWHM at *m*/*z* 200) were averaged to obtain a product ion spectrum for each isolation window.

The set of product ion spectra were collected as a single raw file and deconvolved using the UniChrom module in UniDec version 8.0.3.[1] Briefly, product ion mass spectra were combined into 2 min non-overlapping windows and the resulting averaged spectra were independently deconvolved. Modified deconvolution parameters were used: *m*/*z* range of 2,000 – 8,500, bin every 1 Th, intensity threshold 80, no data normalization, charge range $z$ = 5 – 20, mass range 20 – 50 kDa. The charge smooth width was set to -1, which uses a boxcar function for charge state smoothing and was previously found to reduce deconvolution artifacts.[2] A Gaussian function with FWHM = 2 Th was used as the point spread function in deconvolution. The individual deconvolved spectra were summed to afford a composite spectrum.

**Glycoproteomics**

Proteolytic digestion

OPN and glycosidase-treated OPN samples were separated by SDS-PAGE. Gel bands corresponding to OPN were excised, diced, destained by repeated washes with 25 mM ammonium bicarbonate ($NH_4HCO_3$) in 50% (v/v) acetonitrile (ACN) in LC-MS grade water, and dried in a vacuum concentrator (Savant SPD1010 SpeedVac; Thermo Fisher Scientific). Dried gel pieces were subsequently reduced with 10 mM dithiothreitol (DTT; Merck) in 25 mM $NH_4HCO_3$ for 1 h at 56 °C, followed by alkylation with 55 mM iodoacetamide (Merck) in 25 mM $NH_4HCO_3$ for 45 min at 2 °C in the dark. After washing with 25 mM $NH_4HCO_3$, gels pieces were dehydrated with 25 mM

$NH_4HCO_3$ in 50% ACN and dried. Three combinations of double digests were performed for each sample: Endoproteinase GluC (NEB, P8100S) + Chymotrypsin (CT; Promega, V1061), GluC + O-glycoprotease IMPa (NEB, P0761S), and CT + IMPa. Gel pieces were rehydrated on ice with 10 µL of the corresponding enzymes in 25 mM $NH_4HCO_3$ (CT, GluC 25 ng/µL; IMPa: 100 units/mL) for 10 min, then incubated overnight at 37 °C. For IMPa digests, samples were preincubated with IMPa alone for 2.5 h at 37 °C prior to addition of the second protease. The digest supernatant (aqueous extraction) was collected, and peptides were extracted from the gel pieces with 50% ACN/5% formic acid (FA), combined with the aqueous extraction and dried. Samples were reconstituted in 20 µL buffer A (0.1% (v/v) FA in LC-MS grade water) for LC-MS/MS analysis.

LC-MS/MS method

LC-MS/MS analysis was performed on a Dionex UltiMate 3000 nano HPLC system (Thermo Fisher Scientific) coupled to a hybrid quadrupole-linear ion trap-Orbitrap mass spectrometer (Orbitrap Eclipse Tribrid; Thermo Fisher Scientific). Peptides from the in-gel digest were loaded onto a C18 trap column (PepMap Neo Trap Cartridge, 300 µm × 5 mm, C18, 5 µm, 100 Å; Thermo Fisher Scientific) at a flow rate of 10 µL/min, washed with 60 µL buffer A and separated over an analytical C18 column (Acclaim PepMap 100, 75 µm × 15 cm, nanoviper, C18, 3 µm, 100 Å; Thermo Fisher Scientific). Gradient elution was performed over 120 min at a flow rate of 500 nL/min; the gradient was held at 3% buffer B (0.1% FA in 80% ACN/20% LC-MS grade water) for 6 min, increased linearly from 3% at 6 min to 25% buffer B at 86 min, followed by linear increases to 40% buffer B at 102 min and to 95% buffer B at 106 min, isocratic flow at 95% buffer B from 106 min to 116 min, linear decrease to 3% buffer B at 118 min and re-equilibration at 3% buffer B for 2 min. Eluting peptides were ionized from a stainless-steel emitter (Nano Bore; Thermo Fisher Scientific) using a nESI Flex ionization source (Thermo Fisher Scientific) with an applied spray voltage of 2.3 kV.

Mass spectrometry data were acquired using an optimized data-dependent acquisition approach whereby higher-energy collisional dissociation (HCD) scans are followed by product-dependent electron-transfer dissociation with supplemental HCD (EThcD; HCD-pd-EThcD).[3,4] This ensures EThcD spectra are only acquired for bona fide glycopeptides. Survey scans were acquired in the Orbitrap over a *m*/*z* range of 400 – 1,800 with an automatic gain control (AGC) target of 400,000 charges, a maximum injection time of 50 ms, RF Lens of 50% and a resolution of 60,000 FWHM at *m*/*z* 200. Monoisotopic precursors with charge states *z* = 2 – 8 were selected using the quadrupole with an isolation window of *m*/*z* 2 for data-dependent MS/MS scans (3 s cycle time). Priority filters were set to favor the highest precursor charge states and lowest precursor *m*/*z* values and dynamic exclusion was enabled using a repeat count of 2, a repeat duration of 20 s, a mass tolerance of ± 10 ppm and an exclusion duration of 20 s. Selected ions were fragmented with a normalized collision energy (NCE) of 36% and detected in the Orbitrap with an automated scan range determination (first mass 100 Th), an AGC target of 50,000 charges, a maximum injection time of 54 ms, and a resolution of 30,000 FWHM at *m*/*z* 200. A second EThcD MS/MS scan was acquired if the presence of 2 out of 11 oxonium ion masses (*m*/*z* 126.0550, 138.0549, 144.0655, 163.0601, 168.0654, 186.0760, 204.0865, 274.0921, 292.1027, 325.1120, 366.1395) was detected in the initial HCD MS/MS scan with a mass tolerance of ± 20 ppm. These ions had to be among the 20 most intense peaks in the HCD MS/MS spectrum to trigger EThcD scans.

EThcD MS/MS spectra were recorded in the Orbitrap with the following parameters: 1 s cycle time, an isolation window of *m*/*z* 2, use calibrated charge dependent ETD parameters (to calculate reagent AGC targets and ion-ion reaction times), supplemental HCD activation of 25% NCE, a *m*/*z* range of 200 – 4,000, a maximum injection time of 400 ms, an AGC target of 100,000 charges and a resolution of 60,000 FWHM at *m*/*z* 200.

Data analysis

Glycoproteomics data were analyzed with FragPipe version 23.0, employing the MSFragger-Glyco search engine together with the O-Pair search module.[3,5–7] Searches were performed against a database containing the human OPN (OPNa, UniProt P10451-1) sequence supplemented with common contaminants. Labile mode MSFragger searches used the in-build parameter mass calibration and optimization tool with precursor and fragment *m*/*z* tolerances of 30 ppm and 20 ppm, respectively. In-gel double digests were analyzed by defining the relevant enzyme combinations in the search parameters, with the following enzymatic cleavage settings applied: GluC with up to three missed cleavages, cleaving C-terminal to D and E except when followed by P; CT with up to two missed cleavages, cleaving C-terminal to F, L, W and Y except when followed by P; and IMPa with up to ten missed cleavages, cleaving N-terminal to (glycosylated) S/T residues. For IMPa, the number of allowed missed cleavages was set to up to ten to account for non-glycosylated S/T residues. The searches used the 12-composition human O-glycan database (N1, N1H1, N1A1, N2H1, N1H1A1, N2H2, N2H1A1, N1H1A2, N2H2A1, N2H2F1A1, N2H2A2, N2H2F1A2 – with N for HexNAc, H for Hex, A for NeuAc and F for Fucose) and glycans were defined as mass offsets restricted to S and T residues. Up to four glycosylation sites were allowed per peptide. Oxonium ion filtering was enabled using default diagnostic fragment ions and settings.[7] Other variable modifications that were included in the search were oxidation (M, max 3), protein N-terminal acetylation, phosphorylation (S, T, or Y, max 3) and deamidation (N or Q, max 1). The internal deisotoping and neutral-loss filtering functions were also enabled during database searching. FDR filtering at the PSM, peptide, and protein levels was performed within Philosopher using the PeptideProphet and ProteinProphet modules. O-Pair localization using the O-Pair search module was used for glycosite identification and run for all searches. HCD was set as the first activation type and ETD as the second activation type as described previously.[7] Label-free quantification was performed using MSFragger's IonQuant module, which extracts MS1 precursor intensities.

In-house written scripts were used to aggregate modified peptide forms and compute each form's fractional contribution to the corresponding peptide's total signal based on the MS1 precursor intensities.

**Phosphoproteomics and sulfoproteomics**

Proteolytic digestion and LC-MS/MS method

Concentrated OPN (10 µg) was diluted to 200 µL in 100 mM $NH_4HCO_3$, pH 8.0 (~20-fold dilution) before reduction with DTT and alkylation with iodoacetamide as previously described[8], and digestion with 50:1 (w/w) trypsin gold (Promega) for 18 h with 600 rpm shaking at 37 °C. Digests were then subjected to strong cation exchange using in-house packed stage tip clean-up and dried by vacuum centrifugation.[9] Dried peptides were solubilized in 20 µL of 3% (v/v) ACN /0.1% (v/v) TFA in water, sonicated for 10 min, and centrifuged at 13,000 × *g* for 15 min at 4 °C before separation and analysis by LC-MS/MS using a Dionex UltiMate 3000 nano HPLC system (Thermo Fisher Scientific), over a 30-min gradient, as previously described.[8]

Briefly, samples were loaded onto a trapping column (PepMap100, 300 µm × 5 mm, C18) in loading buffer (3% (v/v) ACN/0.1% (v/v) TFA) at a flow rate of 12 µL/min before being resolved on an analytical column (Easy-Spray C18, 75 µm × 500 mm, 2 µm bead diameter column) using a gradient of 97% A (0.1% (v/v) FA): 2% B (100% (v/v) ACN, 0.1% (v/v) FA) to 40% B over 30 min at a flow rate of 300 nL/min. All data acquisition was performed using a Thermo Orbitrap Fusion Lumos Tribrid mass spectrometer (Thermo Fisher Scientific). MS settings: charge states $z = 2 – 5$, 3 s cycle time, MS1 spectra were acquired in the Orbitrap (60,000 resolution FWHM at $m/z$ 200) over a $m/z$ range of 350–2,000, AGC target = 50%, maximum injection time = 100 ms, with an intensity threshold for fragmentation of 5e5. MS2 spectra were acquired in the Orbitrap (50,000 resolution FWHM at $m/z$ 200), AGC target = standard, maximum injection time = dynamic. A dynamic exclusion window of 5 s was applied at a 10 ppm mass tolerance. For HCDall methods, HCD fragmentation was set at 32% normalized collision energy (NCE). For neutral-loss-triggering methods, initial HCD fragmentation was set at 10% NCE, MS2 acquired in the Orbitrap (15,000 resolution FWHM at $m/z$ 200) and targeted loss continuation trigger set for $m/z$ values equivalent to 1 – 3 sites of sulfation (79.9568 amu) at charge states $z = 2 – 5$ (10 ppm mass tolerance). Continuation of the trigger was performed only for ions that generated NL ion products with at least 10% relative intensity for correct charge state-assigned losses. Triggered scans were acquired in the Orbitrap (50,000 resolution FWHM at $m/z$ 200), HCD set to 32% NCE, AGC target = standard and maximum injection time = auto.

Data analysis

Data were analyzed by Proteome Discoverer 2.4 using the UniProt human database (updated March 2026), searched with fixed modification = carbamidomethylation (C), variable modifications = oxidation (M), deamidation (NQ), phospho (S/T/Y) and sulfo (S/T/Y), instrument type = electrospray ionization-Fourier-transform ion cyclotron resonance (ESI-FTICR), MS1 mass tolerance = 10 ppm, MS2 mass tolerance = 0.01 Da, the Minora Feature Detector node on for label-free quantification (LFQ), and the ptmRS node on. For neutral-loss-triggered data the scan event filter node was included, set to: activation type = HCD, min collision energy = 30, max collision energy = 40. For minimum occupancy calculations, the LFQ intensity of the modified (phospho S/T/Y or sulfo S/T/Y) peptide was divided by the sum of the LFQ intensities of the modified and corresponding unmodified peptide. Note that phospho- or sulfopeptides ionize less efficiently than their unmodified counterparts,[10,11] meaning that occupancy estimates calculated from the ratio of modified peptide to total peptide intensity likely represent minimum values. In instances where the same site modification is observed in multiple peptide forms, the form with the highest Mascot ion score was selected.

**Forward compositional simulation of intact OPN mass distributions**

Intact OPN mass distributions were reconstructed by convolving an underlying protein mass distribution with per-glycan mass distributions representing candidate glycosylation models.

The underlying protein mass distribution, *d*, was derived from the nMS spectrum of OPN FL N1H1 corresponding to the proteoform carrying five O-glycans, by subtracting the mass contribution of five core-1 O-glycans. This distribution therefore retains any residual +80 Da modifications present. The relative abundances of proteoforms carrying five, six, or seven O-glycans, $p_N$, were determined from the simplified mass spectrum of LPP-treated OPN FL N1H1. Per-glycan mass distributions, $g$, representing alternative glycosylation models were constructed from the experimentally determined glycoproteomic data. Because the number of glycans attached to each protein molecule, $N$, varies, the simulated intact mass distribution was modeled as a weighted sum over the three observed occupancy states ($N$ = 5, 6, or 7):

$$\text{Intact OPN mass distribution} = \sum_{N \in \{5,6,7\}} p_N (d * g^{*N})$$

where $g^{*N}$ denotes the $N$-fold self-convolution of the single-glycan mass distribution $g$, representing the combined mass contribution of $N$ stacked glycans.

#### Literature PTMs

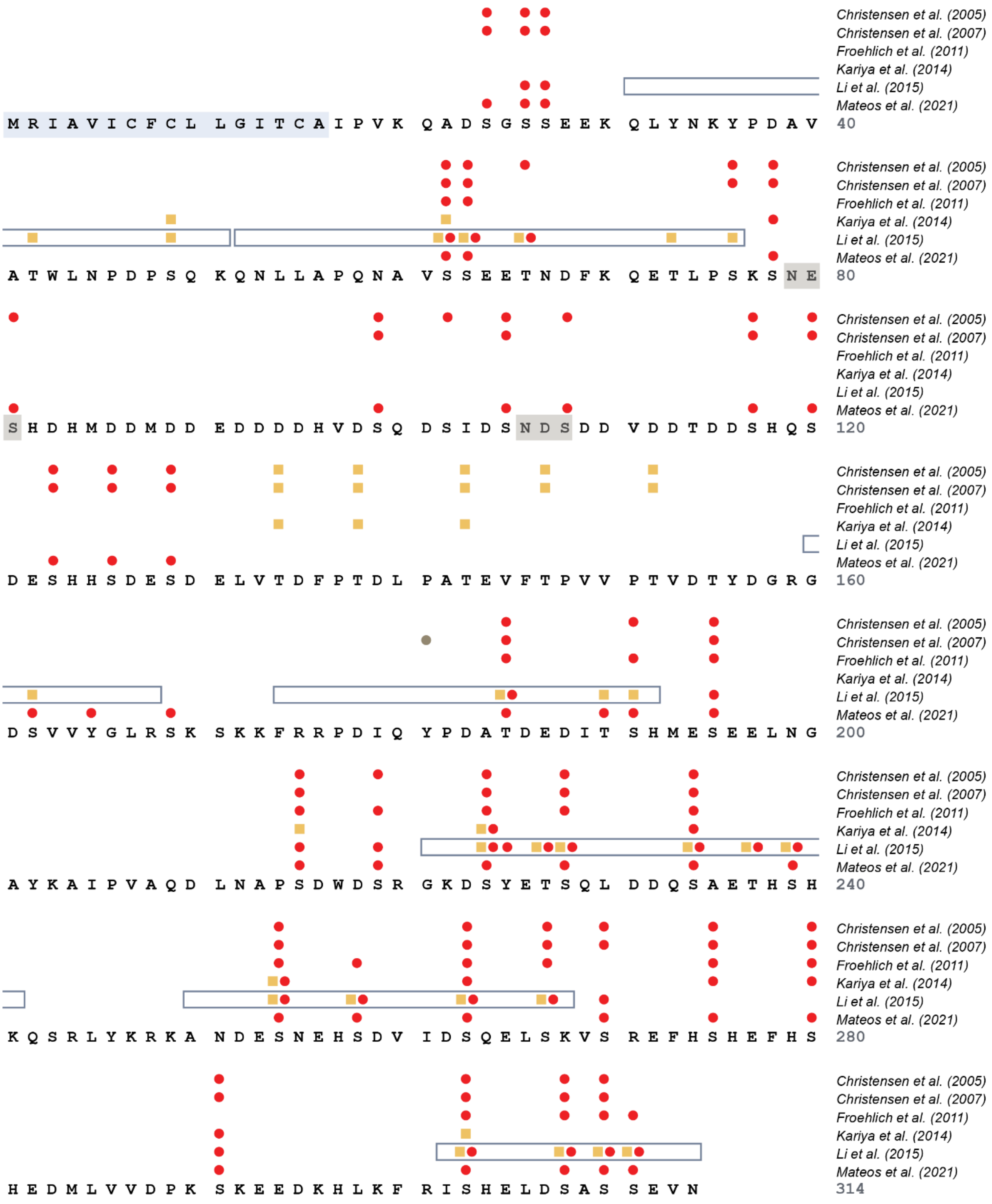

**Figure S1: Post-translational modifications (PTMs) of OPN reported in the literature.** OPN's sequence is heterogeneously and extensively modified. Identified modifications include O-glycosylation (yellow square), phosphorylation (red circle) and sulfation (gray circle). Human OPN originating from milk[12,13], urine[14], or HEK293 cells[15,16], as well as *in vitro* phosphorylated OPN[17], was PTM-profiled. The signal peptide and the N-X-S/T motif required for N-glycosylation are highlighted in light blue and gray, respectively. Boxes indicate glycosylated regions reported in[16].

### DIA-PTCR MS/MS of OPN

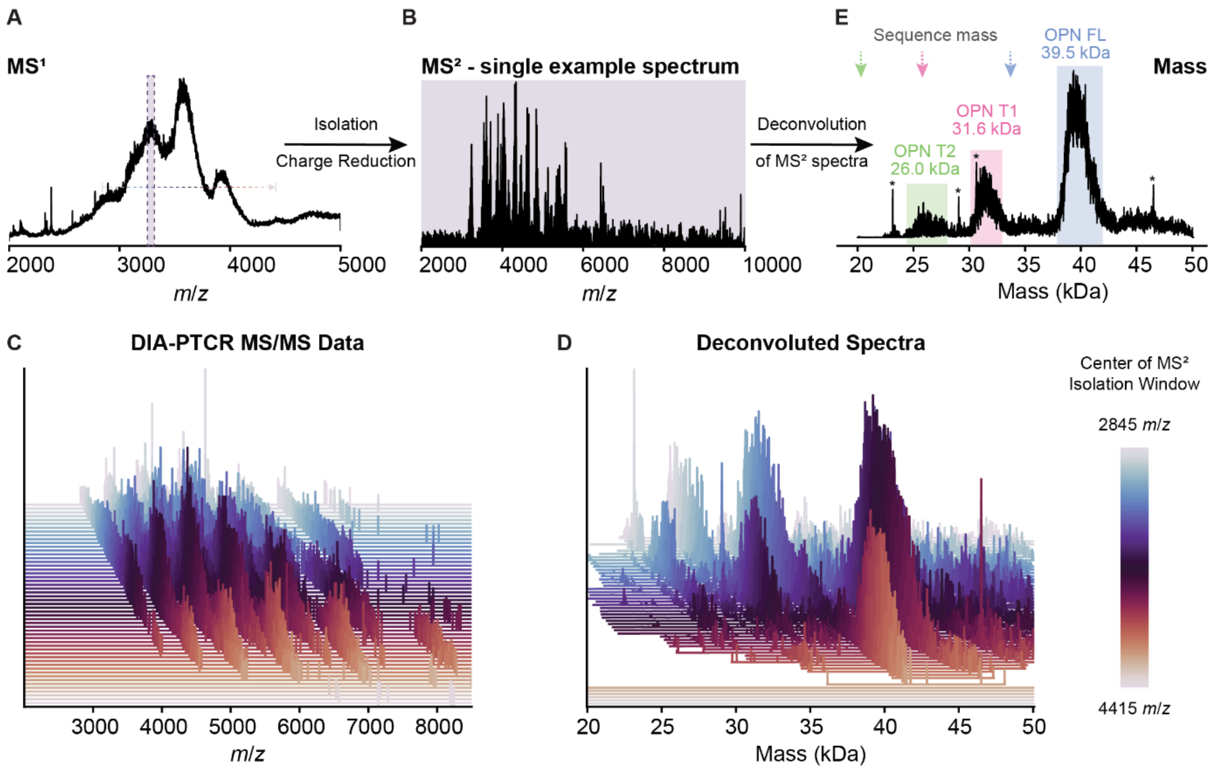

**Figure S2: Capturing the distribution of the OPN proteoforms with data-independent (DIA) proton-transfer charge-reduction (PTCR). (A)** Narrow *m/z* windows (*m/z* 15) in *m/z* range of 2,845 – 4,415 were sequentially isolated from the MS$^1$ spectrum of OPN (1 M ammonium acetate, pH 7.0) and subjected to charge reduction. **(B)** Simplified MS$^2$ spectrum obtained by isolation and charge reduction of a subset of ions. **(C)** Stack of charge-reduced DIA-PTCR MS$^2$ spectra generated by sequential isolation and charge reduction. **(D)** Stack of deconvolved mass spectra resulting from deconvolution of the individual MS$^2$ spectra. **(E)** The combined deconvolved mass spectrum contains three broad peaks corresponding to glycosylated full-length OPN (FL) and its truncated forms (OPN T1 and OPN T2). Arrows mark the sequence-predicted masses of 33.7 kDa, 25.8 kDa, and 20.3 kDa for OPN FL, OPN T1, and OPN T2, respectively.

#### Enzymatic glycan trimming

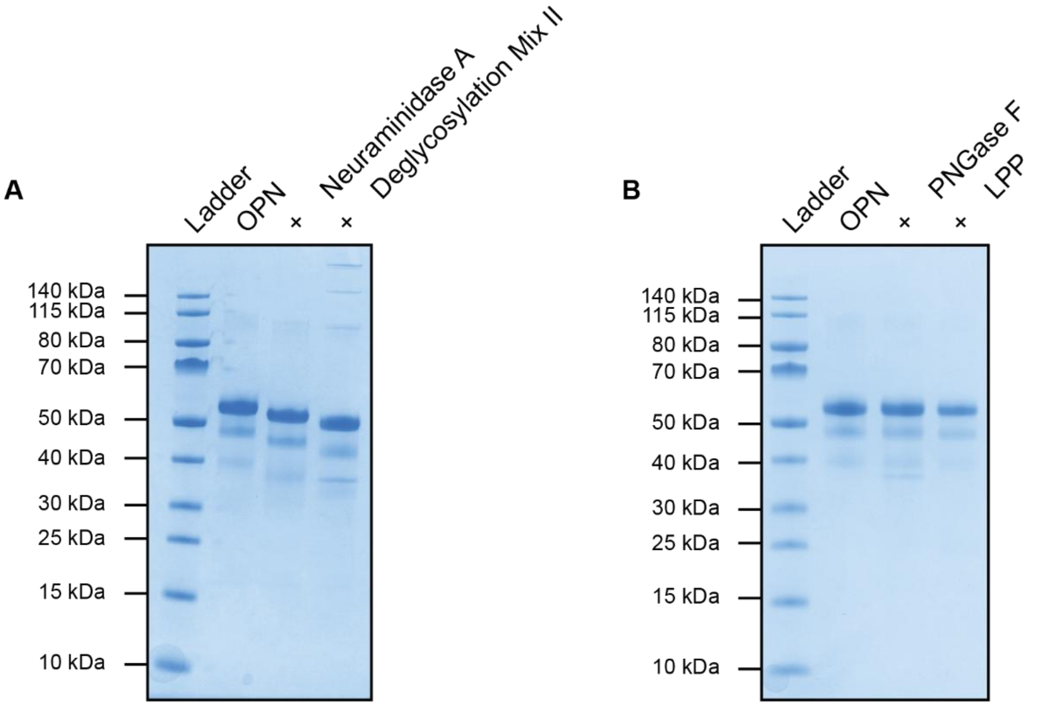

**Figure S3: SDS-PAGE analysis of human OPN and its deglycosylated or dephosphorylated versions. (A)** Untreated OPN shows three bands corresponding to full-length OPN and two truncations. Removal of terminal sialic acids (neuraminidase A) and further deglycosylation (deglycosylation mix II) results in a gel shift. **(B)** No significant gel shift was observed after N-glycan removal (PNGase F) or dephosphorylation (lambda protein phosphatase, LPP), indicating that OPN is not N-glycosylated and not extensively phosphorylated.

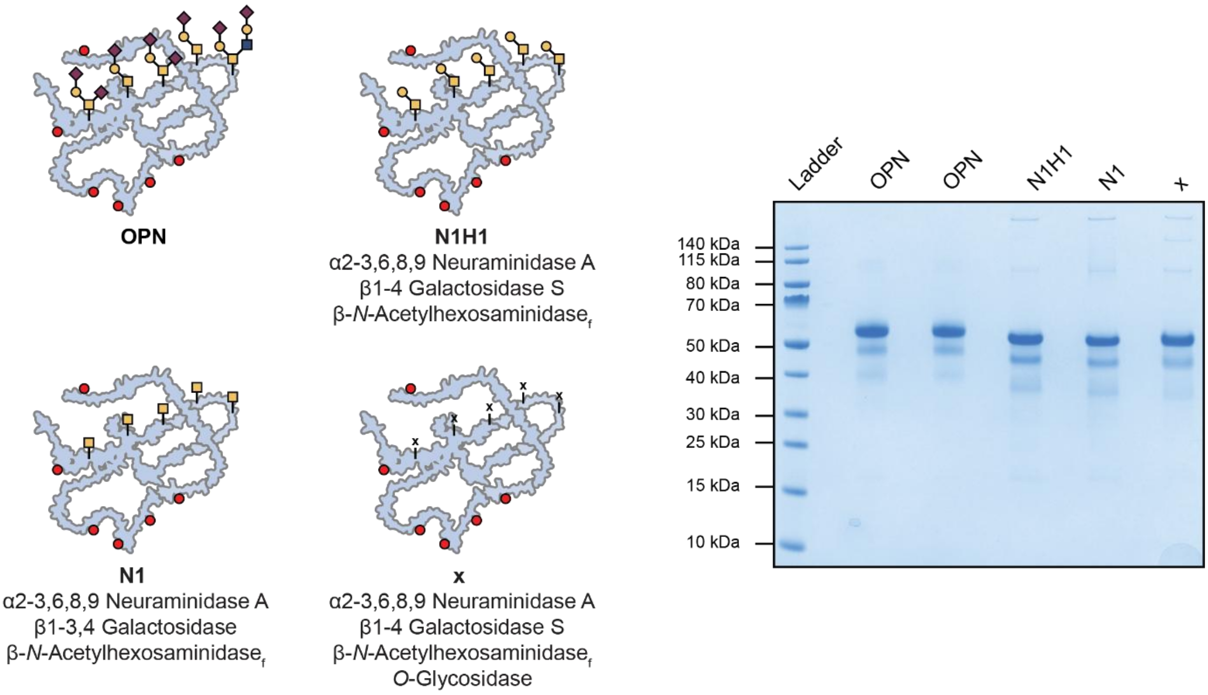

**Figure S4: Enzymatic glycan trimming reduces heterogeneity and creates distinct OPN glycoforms.** Enzymatic glycan removal using *α*2-3,6,8,9 neuraminidase A, *β*1-4 galactosidase S and *β*-*N*-acetylhexosaminidase$_f$ (combination N1H1) transforms (extended) core-1 and core-2 O-glycans into core-1 glycans. Substituting *β*1-3,4 galactosidase for *β*1-4 galactosidase S (combination N1) yields a single GalNAc residue. A combination of *α*2-3,6,8,9 neuraminidase A, *β*1-4 galactosidase S, *β*-*N*-acetylhexosaminidase$_f$ and *O*-glycosidase (combination x) yields full deglycosylation. SDS-PAGE confirms that these treatments alter OPN's apparent molecular weight, but lacks the resolution to resolve the underlying proteoforms or glycan compositions.

#### Phosphorylation state

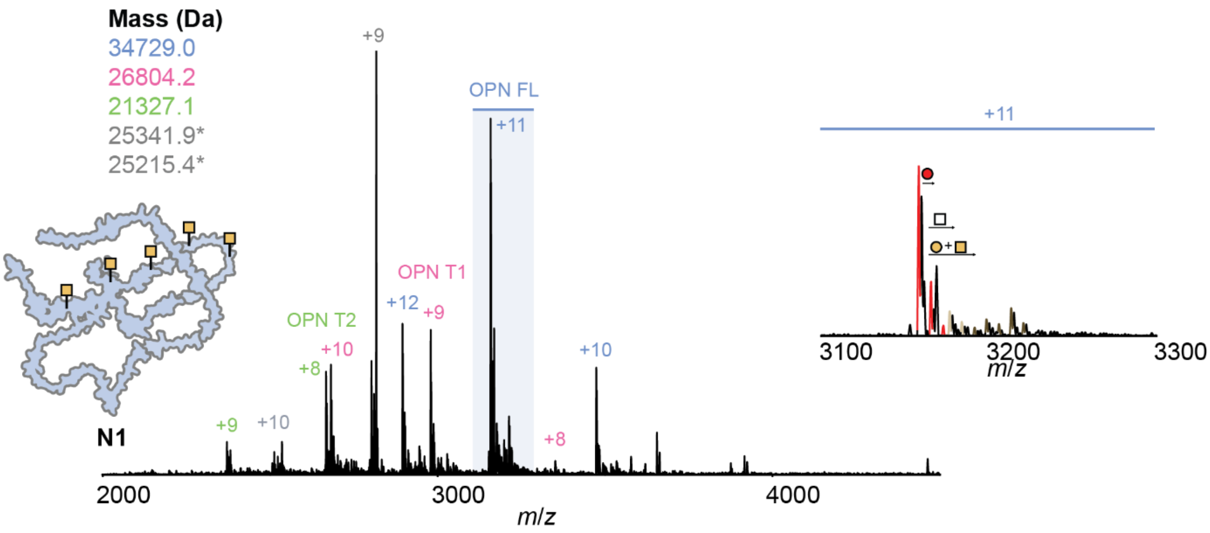

**Figure S5: Native mass spectrum of glycan-trimmed and dephosphorylated OPN N1.** Enzymatic glycan-trimming in combination with dephosphorylation yields a simplified, analyzable mass spectrum (1 M ammonium acetate, pH 7.0) of OPN. Peaks correspond to full-length OPN (FL) and two truncation variants (OPN T1 and OPN T2). All three OPN versions predominantly carry five residual GalNAc residues. The magnification of charge state +11 (inset) highlights residual +80 Da modifications and low occupancy glycosylations. Masses marked with * correspond to lambda protein phosphatase (LPP) used for dephosphorylation.

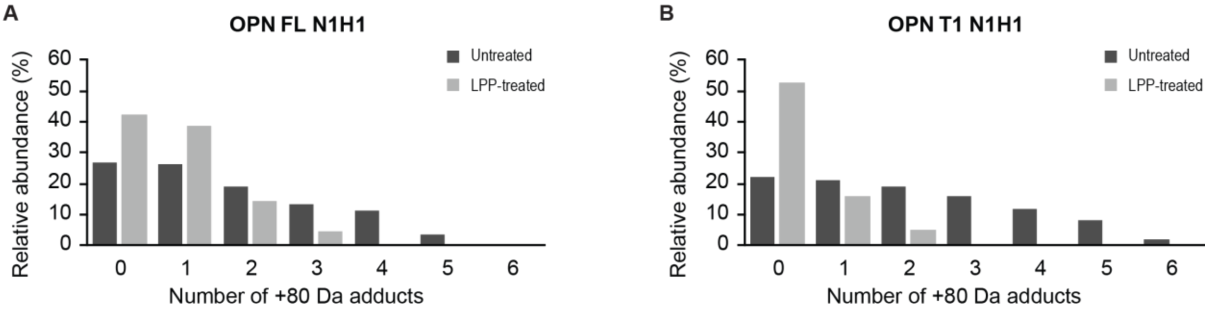

**Figure S6: Comparison of the +80 Da modification-state distribution of intact OPN before and after LPP treatment.** Both **(A)** OPN FL N1H1 and **(B)** OPN T1 N1H1 show a shift toward fewer +80 Da modifications following dephosphorylation, consistent with the removal of phosphate groups. The data were derived from the native mass spectra of OPN N1H1 and dephosphorylated OPN N1H1.

**Table S1: Phosphorylation sites reported for human OPN (P10451) in iPTMnet[18] with a confidence score ≥2, compared against sites covered in the phosphoproteomics dataset, which covered 28 of 41 (68%) reported sites.**

| Site | PTM Type | Score | Within Coverage |
|---|---|---|---|
| S24 | Phosphorylation | 3 | Yes |
| S26 | Phosphorylation | 4 | Yes |
| S27 | Phosphorylation | 4 | Yes |
| S62 | Phosphorylation | 4 | Yes |
| S63 | Phosphorylation | 4 | Yes |
| T66 | Phosphorylation | 3 | Yes |
| S76 | Phosphorylation | 3 | Yes |
| S81 | Phosphorylation | 4 | No |
| S99 | Phosphorylation | 3 | No |
| S102 | Phosphorylation | 3 | No |
| S105 | Phosphorylation | 3 | No |
| S108 | Phosphorylation | 3 | No |
| S117 | Phosphorylation | 3 | No |
| S120 | Phosphorylation | 3 | No |
| S123 | Phosphorylation | 3 | No |
| S126 | Phosphorylation | 3 | No |
| S129 | Phosphorylation | 3 | No |
| T185 | Phosphorylation | 3 | Yes |
| T190 | Phosphorylation | 2 | Yes |
| S191 | Phosphorylation | 4 | Yes |
| S195 | Phosphorylation | 4 | Yes |
| S215 | Phosphorylation | 4 | No |
| S219 | Phosphorylation | 4 | No |
| S224 | Phosphorylation | 4 | Yes |
| Y225 | Phosphorylation | 2 | Yes |
| S228 | Phosphorylation | 4 | Yes |
| S234 | Phosphorylation | 4 | Yes |
| T237 | Phosphorylation | 2 | Yes |
| S239 | Phosphorylation | 2 | Yes |
| S254 | Phosphorylation | 4 | Yes |
| S258 | Phosphorylation | 2 | Yes |
| S263 | Phosphorylation | 4 | Yes |
| S267 | Phosphorylation | 4 | Yes |
| S270 | Phosphorylation | 2 | Yes |
| S275 | Phosphorylation | 4 | Yes |
| S280 | Phosphorylation | 4 | Yes |
| S291 | Phosphorylation | 2 | No |
| S303 | Phosphorylation | 4 | Yes |
| S308 | Phosphorylation | 4 | Yes |
| S310 | Phosphorylation | 4 | Yes |
| S311 | Phosphorylation | 2 | Yes |

**Table S2: Overview of the peptides quantified in phosphoproteomics and sulfoproteomics experiments.** Phosphorylation was quantified at 12 sites: Ser26/Ser27, Ser62, Ser63, Ser195, Ser234, Ser254, Ser263, Ser275, Ser280, Ser310, and Ser311. Comparative analysis of untreated and LPP-treated OPN samples using a custom proteomics workflow for the identification of sulfotyrosine-containing peptides[11] identified two candidate sulfotyrosine-containing peptides: R$^{176}$PDIQYPDATDEDITSHMESEELNGAYK$^{203}$ and G$^{221}$KDSYETSQLDDQSAETHSHK$^{241}$.

| Annotated Sequence | Modifications | Residues | Modification Sites | # PSMs | Mascot Ion Score | Top Apex RT [min] | Untreated | LPP-treated | Minimum Occupancy (%) of Untreated Sample |
|---|---|---|---|---|---|---|---|---|---|
| [K].QADSGSSEEKQLYNKYPDAVATWLNPDPSQK.[Q] | 1xPhospho [S/T] | 21-51 | Unlocalized (S26/S27) | 9 | 45 | 31.86 | 3.22E+06 | | 7.10 |
| [K].QNLLAPQNAVSSEETNDFKQETLPSK.[S] | 1xPhospho [S11(100)] | 52-77 | S62(100) | 27 | 73 | 28.86 | 2.53E+08 | | 17.29 |
| [K].QNLLAPQNAVSSEETNDFKQETLPSK.[S] | 2xPhospho [S11(100); S12(100)] | 52-77 | S62(100); S63(100) | 4 | 54 | 30.85 | 2.22E+07 | | 1.52 |
| [K].FRRPDIQYPDATDEDITSHMESEELNGAYK.[A] | 1xDeamidated [N26]; 1xSulfo [T/Y/S]; 1xPhospho [S22(100)] | 174-203 | S195(100); Unlocalized | 40 | 62 | 29.94 | 1.57E+07 | | 3.06 |
| [R].RPDIQYPDATDEDITSHMESEELNGAYK.[A] | 1xDeamidated [N24]; 1xSulfo [T/S/Y] | 176-203 | Unlocalized | 828 | 88 | 28.98 | 4.18E+08 | 5.73E+08 | 13.01 |
| [R].RPDIQYPDATDEDITSHMESEELNGAYK.[A] | 1xPhospho [S20(100)] | 176-203 | S195(100) | 3 | 56 | 28.49 | 4.56E+08 | 4.87E+07 | 36.54 |
| [R].GKDSYETSQLDDQSAETHSHK.[Q] | 1xSulfo [Y/T/S] | 221-241 | Unlocalized | 238 | 113 | 16.27 | 6.79E+08 | 3.05E+07 | 5.29 |
| [R].GKDSYETSQLDDQSAETHSHK.[Q] | 2xPhospho [S14(100); S/T] | 221-241 | S234(100); Unlocalized | 11 | 75 | 17.44 | 3.66E+07 | | 0.28 |
| [K].DSYETSQLDDQSAETHSHK.[Q] | 1xPhospho [S12(100)] | 223-241 | S234(100) | 10 | 105 | 18.25 | 5.12E+07 | | 3.86 |
| [R].KANDESNEHSDVIDSQELSK.[V] | 1xDeamidated [N]; 1xPhospho [S6(100)] | 249-268 | S254(100) | 23 | 113 | 20.26 | 5.83E+07 | | 2.99 |
| [R].KANDESNEHSDVIDSQELSK.[V] | 2xPhospho [S6(100); S15(100)] | 249-268 | S254(100); S263(100) | 6 | 98 | 20.88 | 3.58E+07 | | 0.48 |
| [K].VSREFHSHEFHSHEDMLVVDPK.[S] | 2xPhospho [S7(100); S12(100)] | 269-290 | S275(100); S280(100) | 3 | 29 | 22.08 | 1.46E+06 | | 6.67 |
| [R].EFHSHEFHSHEDMLVVDPK.[S] | 1xPhospho [S9(100)] | 272-290 | S280(100) | 42 | 69 | 22.35 | 1.34E+08 | 1.07E+05 | 9.64 |
| [K].FRISHELDSASSEVN.[-] | 1xPhospho [S12(99.9)] | 300-314 | S311(99.9) | 49 | 93 | 24.33 | 2.89E+08 | 1.41E+05 | 17.35 |
| [K].FRISHELDSASSEVN.[-] | 2xPhospho [S11(99.8); S12(99.8)] | 300-314 | S310(99.8); S311(99.8) | 8 | 65 | 27.12 | 9.36E+06 | | 0.56 |

**Table S3: Approximated minimum occupancies for each modified OPN peptide region identified by phosphoproteomics.**

| Peptide Start | Peptide End | Unmodified | 1 Modification | 2 Modifications |
|---|---|---|---|---|
| 21 | 51 | 0.929 | 0.071 | 0 |
| 52 | 77 | 0.8119 | 0.1729 | 0.0152 |
| 176 | 203 | 0.8 | 0.2 | 0 |
| 221 | 241 | 0.9443 | 0.0529 | 0.0028 |
| 249 | 268 | 0.9653 | 0.0299 | 0.0048 |
| 269 | 290 | 0.9333 | 0 | 0.0667 |
| 300 | 314 | 0.8209 | 0.1735 | 0.0056 |

**Table S4: Predicted distribution of phosphorylations per OPN molecule.** Probability of a molecule carrying 0–6 phosphorylations, obtained by convolving the per-region minimum occupancy distributions across all seven modified peptide regions, assuming independent modification status between regions. Mean stoichiometry = 0.89 per molecule.

| # of Phosphorylations | Sum | Weighted Fraction |
|---|---|---|
| 0 | 0.421 | 0 |
| 1 | 0.353 | 0.353 |
| 2 | 0.158 | 0.316 |
| 3 | 0.0522 | 0.1566 |
| 4 | 0.013 | 0.052 |
| 5 | 0.00229 | 0.01145 |
| 6 | 0.00029 | 0.00174 |
| | **Sum** | 0.89079 |

**Table S5: Distribution of +80 Da modifications derived from native mass spectrometry for the predominant five- and six-O-glycan proteoforms of LPP-treated intact OPN FL N1H1.**

| # of +80 Da Modifications | 5 Glycans | Fraction | 6 Glycans | Fraction | Sum | Fraction | Weighted Fraction |
|---|---|---|---|---|---|---|---|
| 0 | 0.99 | 0.55 | 0.18 | 0.19 | 1.17 | 0.42 | 0.00 |
| 1 | 0.64 | 0.35 | 0.43 | 0.45 | 1.07 | 0.39 | 0.39 |
| 2 | 0.16 | 0.09 | 0.24 | 0.25 | 0.40 | 0.14 | 0.29 |
| 3 | 0.02 | 0.01 | 0.10 | 0.11 | 0.12 | 0.04 | 0.13 |
| | | | | | | **Sum** | 0.81 |

**Table S6: Distribution of +80 Da modifications derived from native mass spectrometry for the predominant five- and six-O-glycan proteoforms of intact OPN FL N1H1.**

| # of +80 Da Modifications | 5 Glycans | Fraction | 6 Glycans | Fraction | Sum | Fraction | Weighted Fraction |
|---|---|---|---|---|---|---|---|
| 0 | 1.00 | 0.34 | 0.17 | 0.12 | 1.17 | 0.27 | 0.00 |
| 1 | 0.72 | 0.25 | 0.42 | 0.29 | 1.14 | 0.26 | 0.26 |
| 2 | 0.50 | 0.17 | 0.33 | 0.23 | 0.83 | 0.19 | 0.38 |
| 3 | 0.35 | 0.12 | 0.23 | 0.16 | 0.58 | 0.13 | 0.40 |
| 4 | 0.28 | 0.10 | 0.20 | 0.14 | 0.48 | 0.11 | 0.44 |
| 5 | 0.06 | 0.02 | 0.10 | 0.07 | 0.16 | 0.04 | 0.18 |
| 6 | 0.00 | 0.00 | 0.00 | 0.00 | 0.00 | 0.00 | 0.00 |
| | | | | | | **Sum** | 1.67 |

**Table S7: Distribution of +80 Da modifications derived from native mass spectrometry for the predominant five- and six-O-glycan proteoforms of LPP-treated intact OPN T1 N1H1.**

| # of +80 Da Modifications | 5 Glycans | Fraction | 6 Glycans | Fraction | Sum | Fraction | Weighted Fraction |
|---|---|---|---|---|---|---|---|
| 0 | 0.58 | 0.67 | 0.08 | 0.21 | 0.66 | 0.53 | 0.00 |
| 1 | 0.24 | 0.28 | 0.20 | 0.53 | 0.44 | 0.16 | 0.16 |
| 2 | 0.05 | 0.06 | 0.09 | 0.24 | 0.14 | 0.05 | 0.10 |
| 3 | 0.00 | 0.00 | 0.01 | 0.03 | 0.01 | 0.00 | 0.01 |
| | | | | | | **Sum** | 0.27 |

**Table S8: Distribution of +80 Da modifications derived from native mass spectrometry for the predominant five- and six-O-glycan proteoforms of intact OPN T1 N1H1.**

| # of +80 Da Modifications | 5 Glycans | Fraction | 6 Glycans | Fraction | Sum | Fraction | Weighted Fraction |
|---|---|---|---|---|---|---|---|
| 0 | 0.68 | 0.28 | 0.12 | 0.10 | 0.80 | 0.22 | 0.00 |
| 1 | 0.49 | 0.20 | 0.27 | 0.23 | 0.77 | 0.21 | 0.21 |
| 2 | 0.41 | 0.17 | 0.29 | 0.24 | 0.69 | 0.19 | 0.38 |
| 3 | 0.38 | 0.15 | 0.20 | 0.17 | 0.58 | 0.16 | 0.48 |
| 4 | 0.25 | 0.10 | 0.18 | 0.15 | 0.43 | 0.12 | 0.47 |
| 5 | 0.17 | 0.07 | 0.12 | 0.10 | 0.29 | 0.08 | 0.40 |
| 6 | 0.06 | 0.02 | 0.02 | 0.02 | 0.08 | 0.02 | 0.13 |
| | | | | | | **Sum** | 2.07 |

### Glycoproteomics

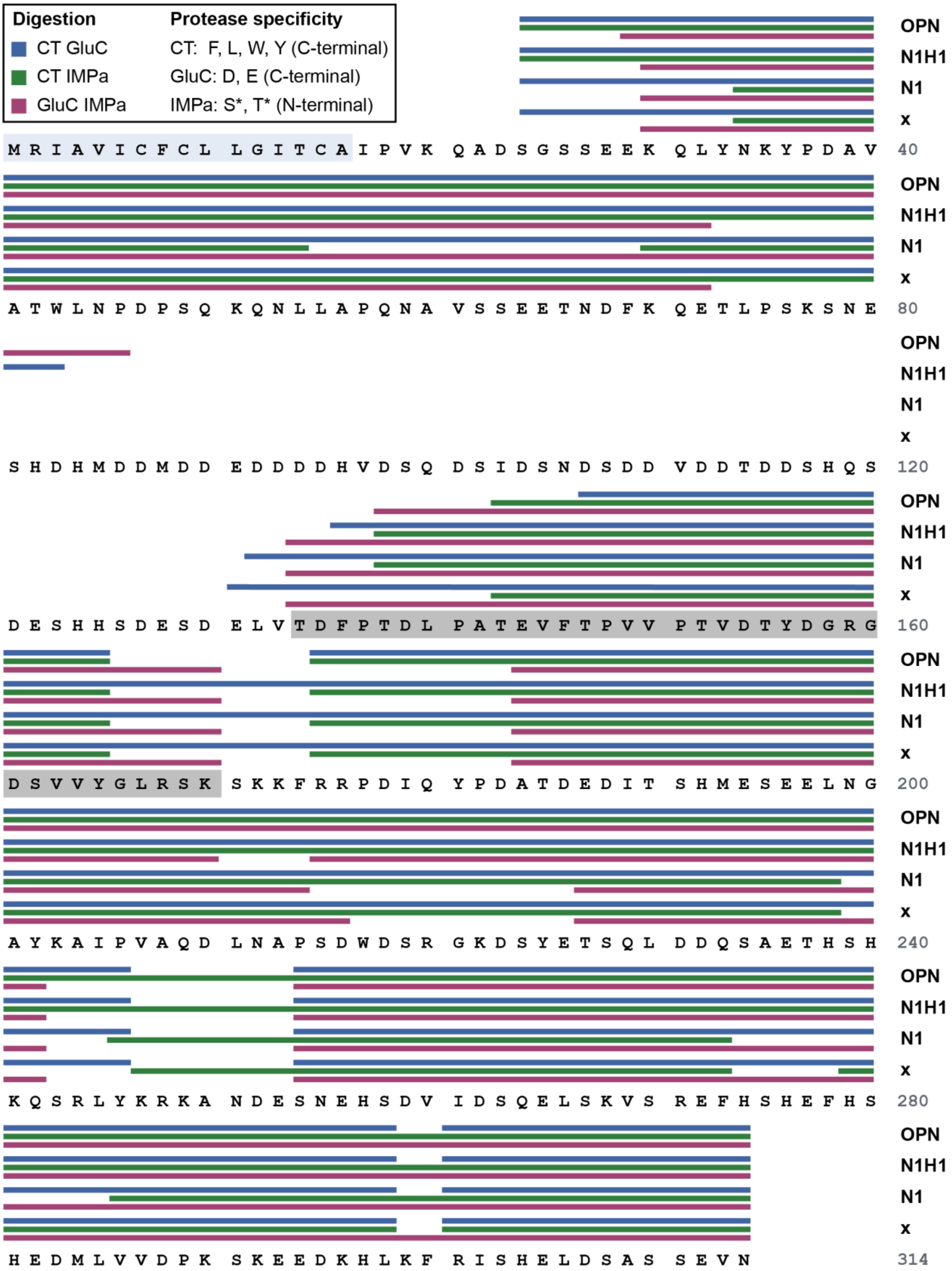

**Figure S7: Sequence coverage of OPN obtained with different protease combinations in the glycoproteomics experiments.** Untreated OPN and exoglycosidase-treated N1H1, N1 and x were digested with combinations of chymotrypsin (CT) and endoproteinase GluC (GluC), CT and O-glycoprotease IMPa as well as GluC and IMPa. CT targets Phe, Tyr, Trp and Leu, while GluC targets Asp and Glu. IMPa is an O-glycan-specific protease, which can cleave peptide bonds immediately N-terminal to glycosylated to Ser/Thr residues, denoted by *. Sequence coverage was obtained across most of the OPN sequence, including the threonine/proline-rich region (gray) that harbors a high density of previously reported glycosites. Coverage was lacking for residues 87 – 130.

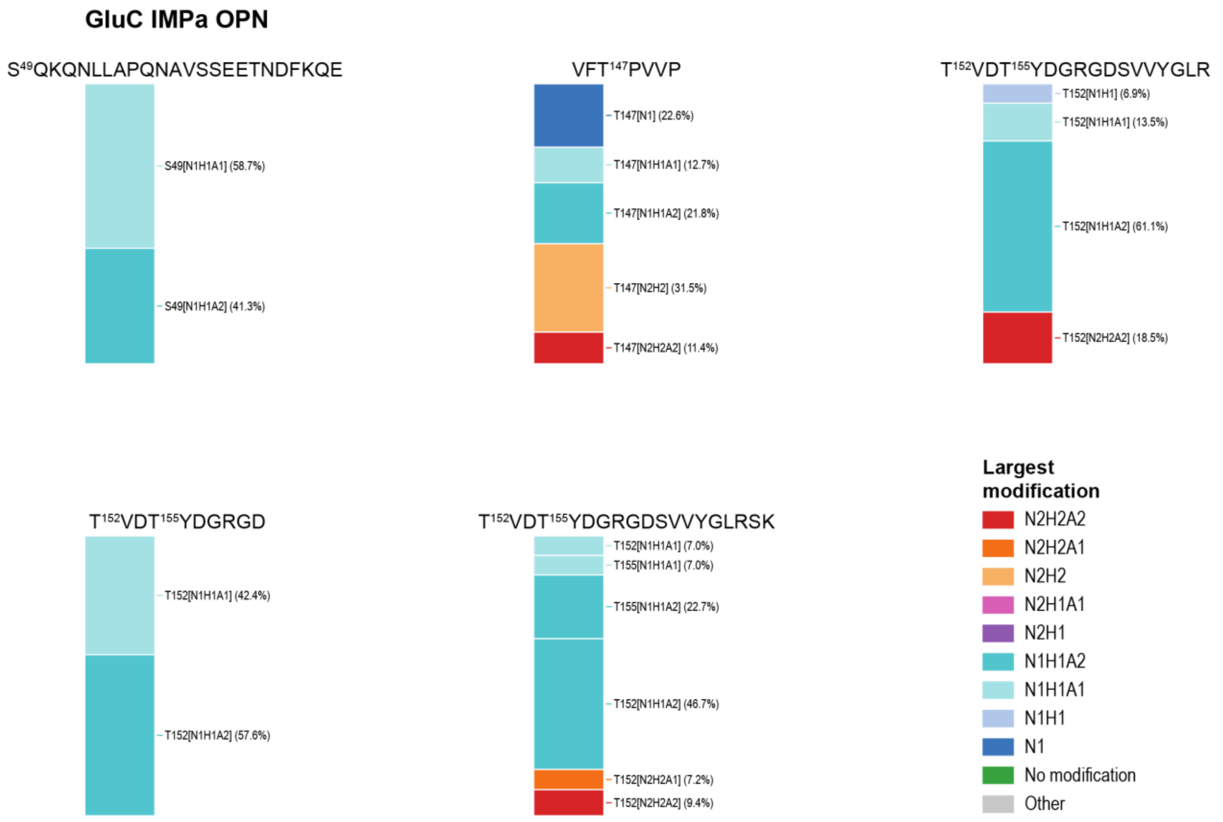

**Figure S8: Glycopeptides observed for untreated OPN digested with GluC and IMPa, excluding peptides already shown in the main text figures.**

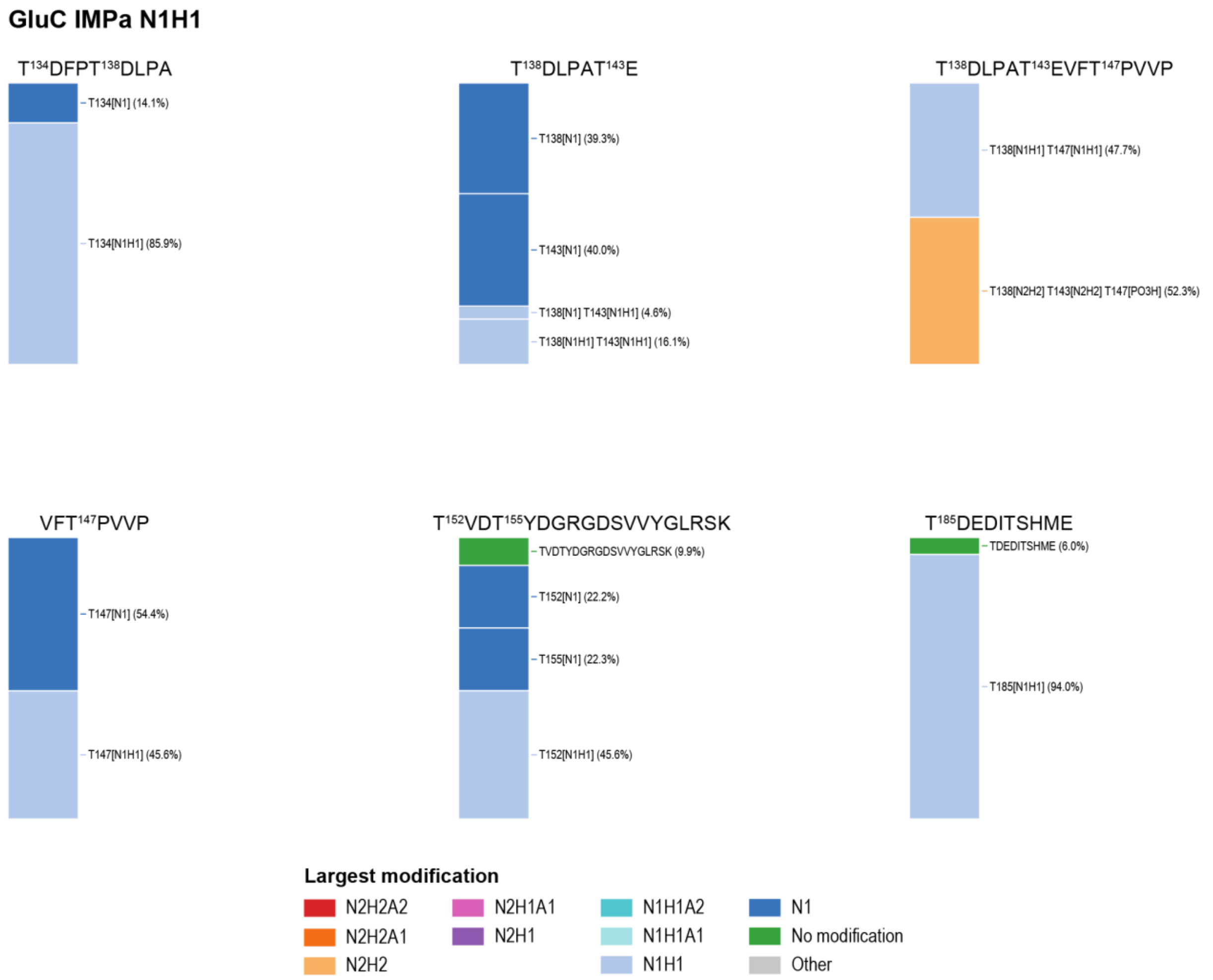

**Figure S9: Glycopeptides observed for exoglycosidase-treated OPN N1H1 digested with GluC and IMPa, excluding peptides already shown in the main text figures.**

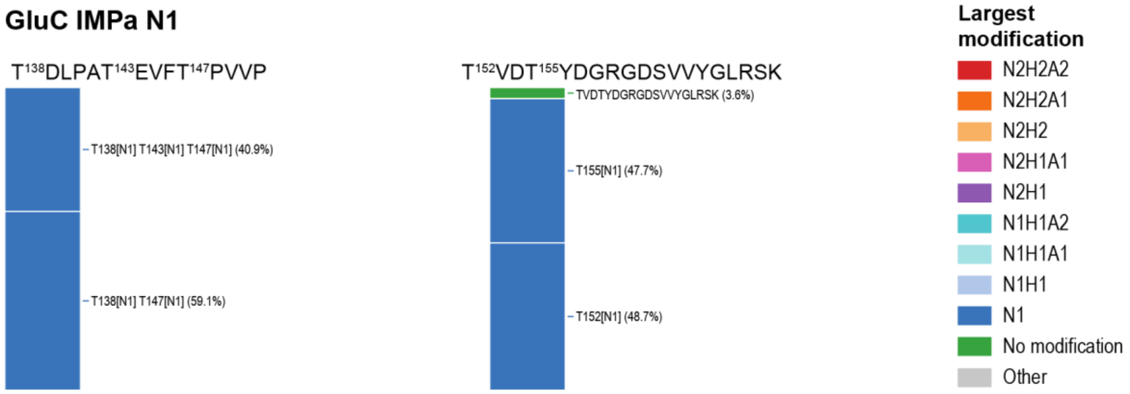

**Figure S10: Glycopeptides observed for exoglycosidase-treated OPN N1 digested with GluC and IMPa, excluding peptides already shown in the main text figures**.

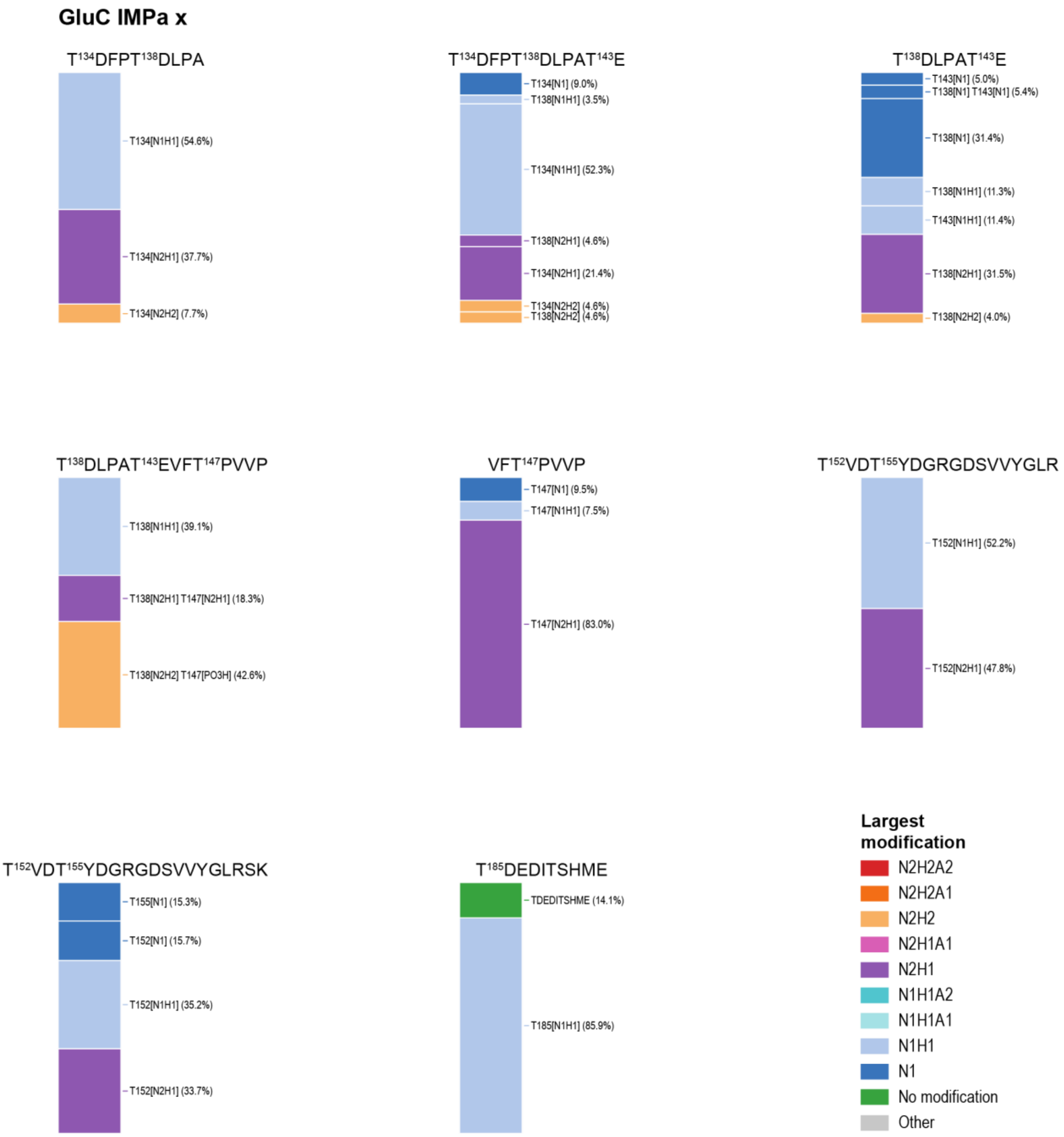

**Figure S11: Glycopeptides observed for deglycosylated OPN x digested with GluC and IMPa, excluding peptides already shown in the main text figures**. In the fully deglycosylated sample, IMPa can no longer cleave at successfully deglycosylated sites, so the peptides observed here primarily originate from sites where glycan release was incomplete.

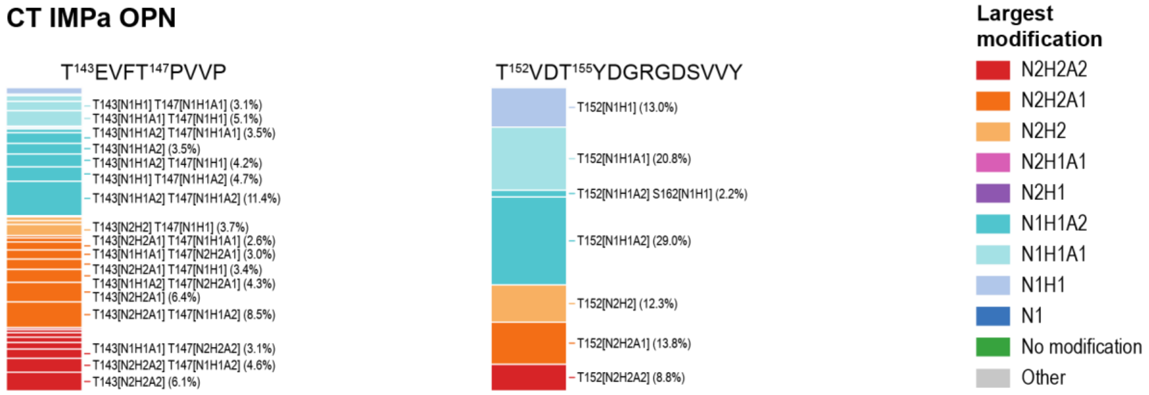

**Figure S12: Glycopeptides observed for untreated OPN digested with CT and IMPa.** Minor species below 2.5% relative abundance are left unannotated where labeling would reduce figure clarity.

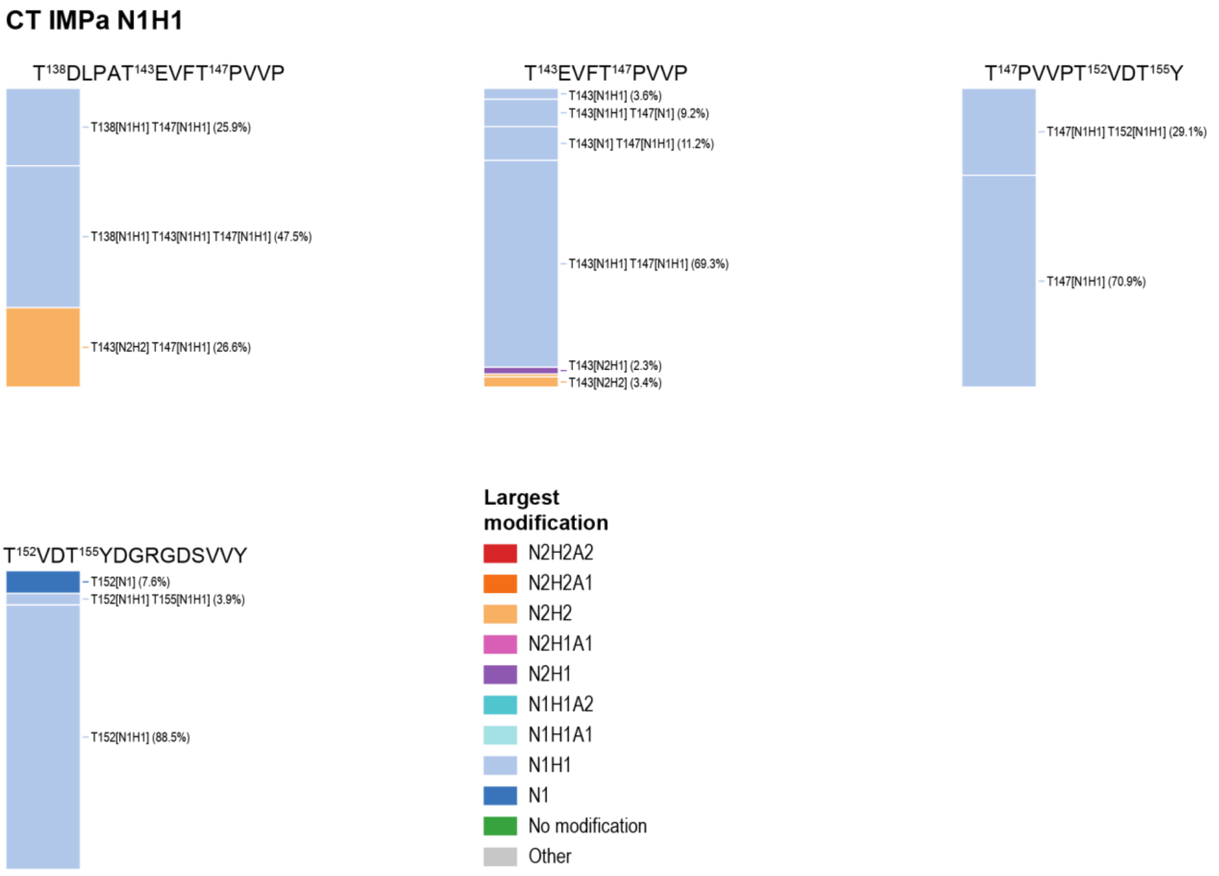

**Figure S13: Glycopeptides observed for exoglycosidase-treated OPN N1H1 digested with CT and IMPa.**

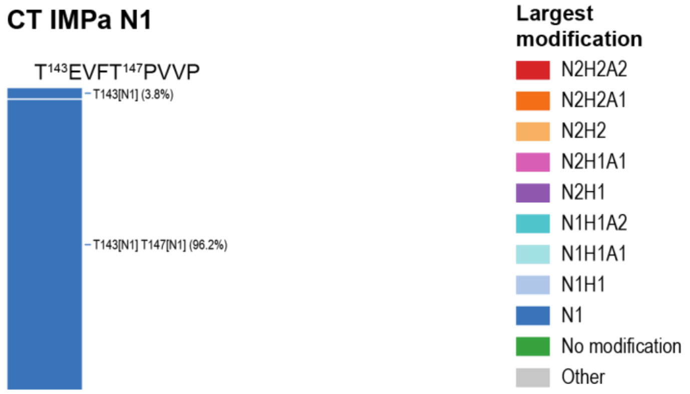

**Figure S14: Glycopeptides observed for exoglycosidase-treated OPN N1 digested with CT and IMPa.**

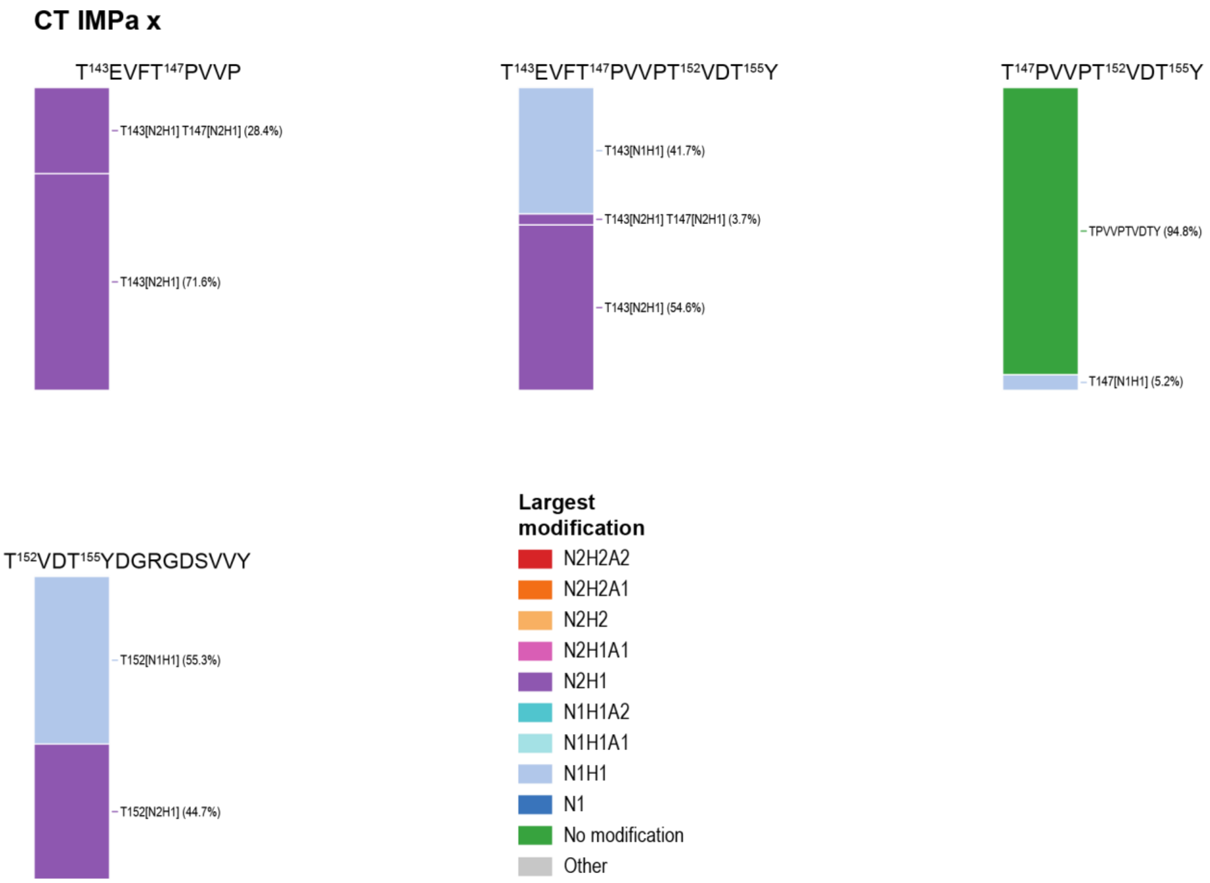

**Figure S15: Glycopeptides observed for deglycosylated OPN x digested with CT and IMPa.** In the fully deglycosylated sample, IMPa can no longer cleave at successfully deglycosylated sites, so the peptides observed here primarily originate from sites where glycan release was incomplete.

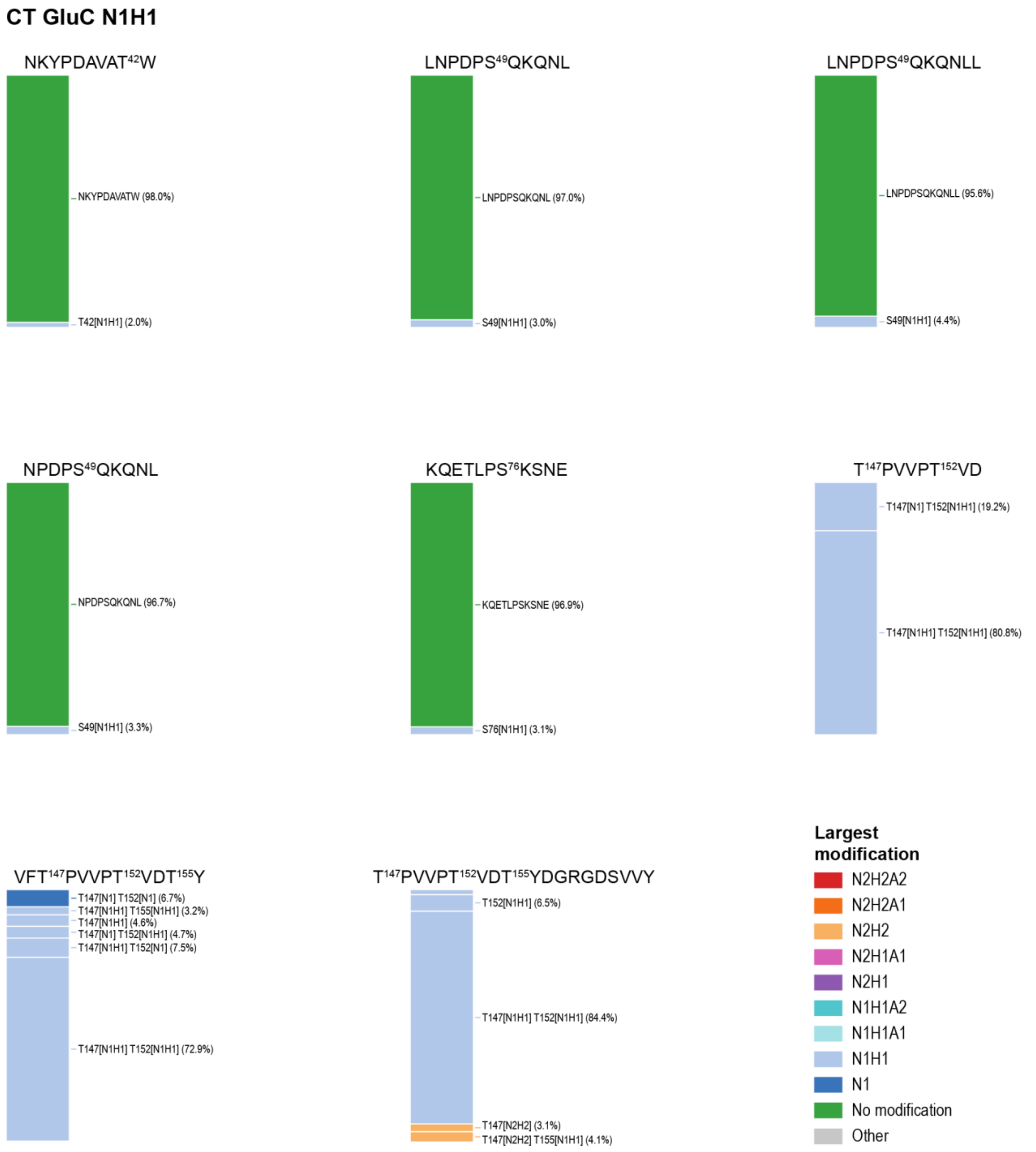

**Figure S16: Glycopeptides observed for exoglycosidase-treated OPN N1H1 digested with CT and GluC, excluding peptides already shown in the main text figures.**

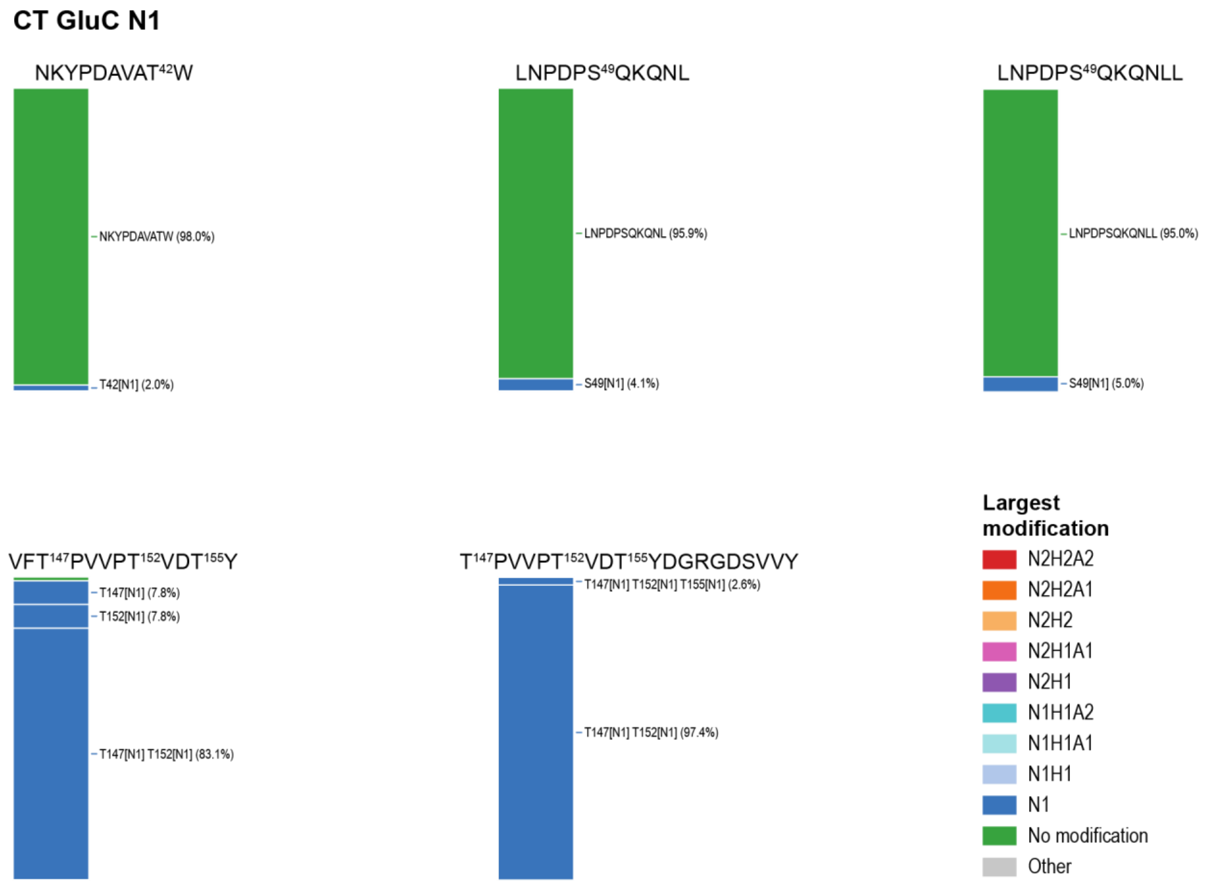

**Figure S17: Glycopeptides observed for exoglycosidase-treated OPN N1 digested with CT and GluC, excluding peptides already shown in the main text figures**. Minor species below 2.5% relative abundance are left unannotated where labeling would reduce figure clarity.

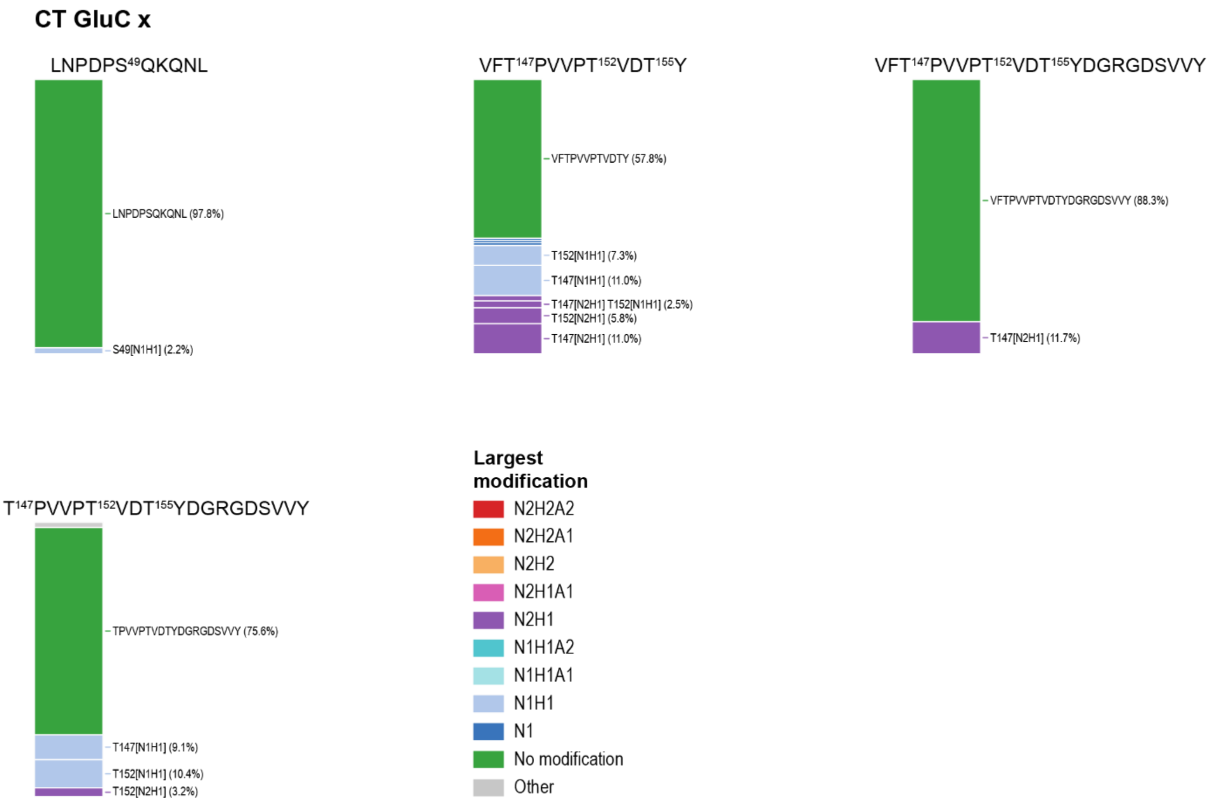

**Figure S18: Glycopeptides observed for deglycosylated OPN x digested with CT and GluC, excluding peptides already shown in the main text figures**. Minor species below 2.5% relative abundance are left unannotated where labeling would reduce figure clarity.

### Simulation of intact OPN mass distributions

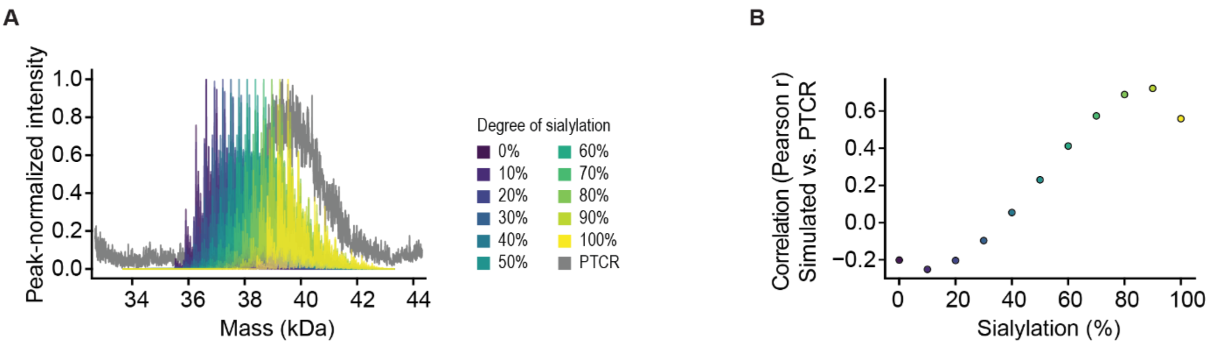

**Figure S19: Titrating glycan sialylation in the forward simulation identifies the sialylation level that best reproduces the experimental intact OPN mass distribution. (A)** Simulated intact OPN mass distributions for per-glycan sialylation fractions ranging from 0% to 100% in 10% increments, with the experimentally determined Thr143 core-1:core-2 ratio (peptide $T^{143}EVFT^{147}PVVP$, GluC/IMPa digestion) held constant, benchmarked against the experimental PTCR mass distribution. **(B)** Pearson correlation coefficient (r) between simulated and experimental PTCR mass distributions as a function of per-glycan sialylation fraction, showing the best agreement at approximately 90% sialylation.

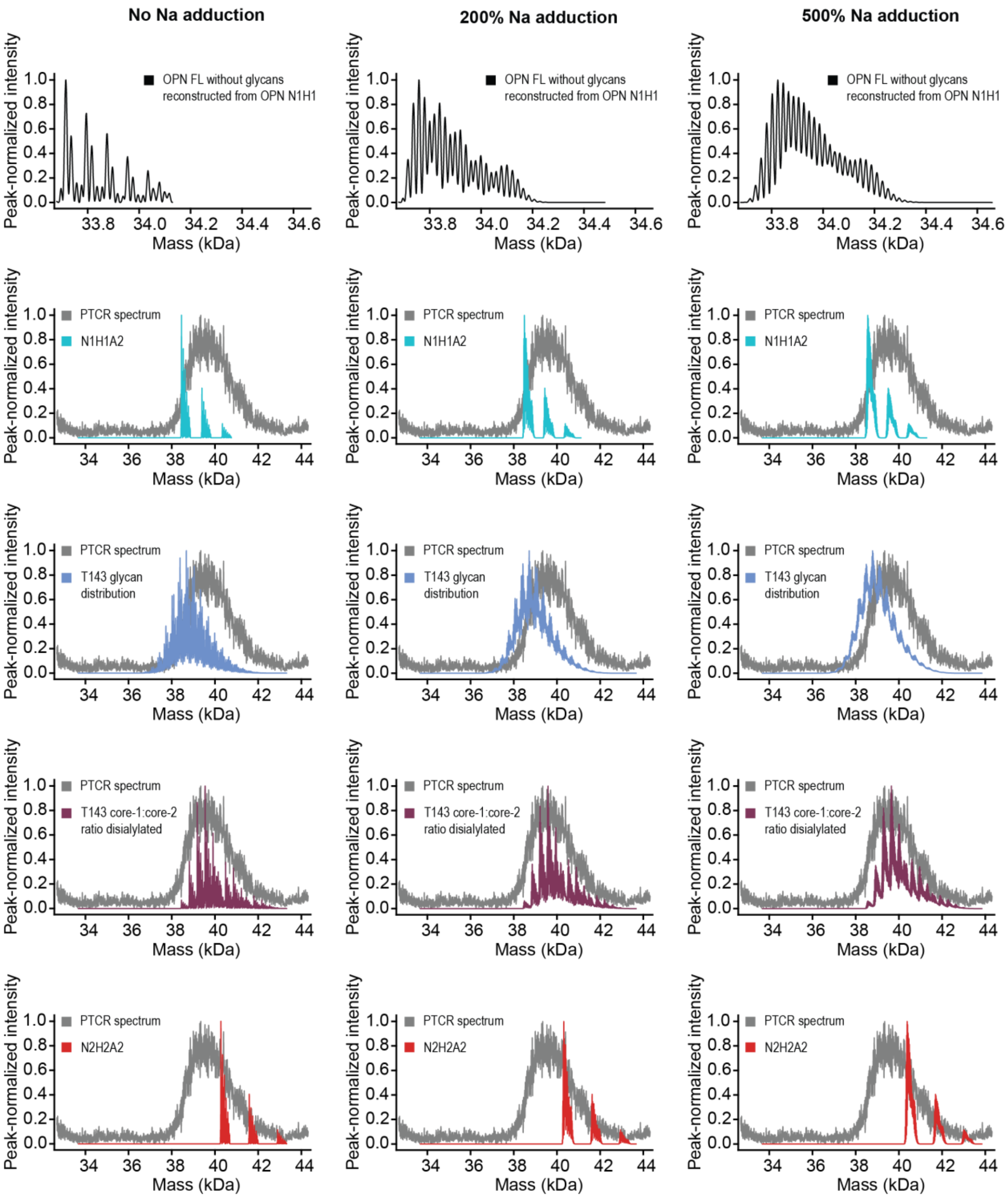

**Figure S20: Sodium adduction broadens simulated peak envelopes but does not change the relative agreement between glycosylation composition models and the experimental PTCR spectrum.** Columns show increasing mean sodium-adduct load per OPN molecule (Poisson adduct ladder, +21.98 Da per $Na^+$ adduct): no adduction (left), 200% mean adduction (middle), and 500% mean adduction (right). Simulated intact OPN FL mass distributions under four glycosylation models: a homogeneous core-1 disialylated glycan at every occupied site (N1H1A2, cyan), the experimental Thr143 glycan compositional profile (T143 glycan distribution, blue), a binary core-1/core-2 disialylated mixture (N1H1A2/N2H2A2) at the experimentally determined Thr143 core-1:core-2 ratio (maroon), and a homogeneous core-2 disialylated glycan at every occupied site (N2H2A2, red).

**Table S9: Phosphorylation and sulfation (+80 Da modifications) distribution and O-glycosylation state (five, six, or seven O-glycans) derived from native mass spectrometry of intact OPN FL N1H1.**

| # of +80 Da Modifications | 5 Glycans | Fraction | 6 Glycans | Fraction | 7 Glycans | Fraction | Sum |
|---|---|---|---|---|---|---|---|
| 0 | 1.00 | 0.21 | 0.17 | 0.03 | 0.07 | 0.01 | 0.25 |
| 1 | 0.72 | 0.15 | 0.42 | 0.09 | 0.10 | 0.02 | 0.25 |
| 2 | 0.50 | 0.10 | 0.33 | 0.07 | 0.10 | 0.02 | 0.19 |
| 3 | 0.35 | 0.07 | 0.23 | 0.05 | 0.10 | 0.02 | 0.14 |
| 4 | 0.28 | 0.06 | 0.20 | 0.04 | 0.06 | 0.01 | 0.11 |
| 5 | 0.06 | 0.01 | 0.10 | 0.02 | 0.07 | 0.01 | 0.05 |
| 6 | 0.00 | 0.00 | 0.00 | 0.00 | 0.03 | 0.01 | 0.01 |
| | **Sum** | 0.60 | **Sum** | 0.30 | **Sum** | 0.11 | |

**Table S10: Relative abundances of proteoforms carrying five, six, or seven O-glycans determined from the simplified mass spectrum of LPP-treated OPN FL N1H1.**

| Total # of Glycans | Relative Abundance | Fraction |
|---|---|---|
| 5 | 100 | 0.66 |
| 6 | 40.6 | 0.27 |
| 7 | 11.6 | 0.08 |

**Table S11: Per-glycan composition distribution and core-1:core-2 ratio for Thr143, derived from glycoproteomic analysis of peptide T[143]EVFT[147]PVVP following GluC and IMPa digestion of untreated OPN.**

| Glycan type | Mass (Da) | Fraction of T143 |
|---|---|---|
| N1 | 203.0794 | 0.00 |
| N1H1 | 365.1322 | 0.04 |
| N2H1 | 568.2116 | 0.00 |
| N2H2 | 730.2644 | 0.08 |
| N1H1A1 | 656.2276 | 0.21 |
| N2H1A1 | 859.307 | 0.00 |
| N1H1A2 | 947.323 | 0.24 |
| N2H2A1 | 1021.3598 | 0.22 |
| N2H2A2 | 1312.4552 | 0.22 |
| Core-1 | | 0.48 |
| Core-2 | | 0.52 |

**Table S12: Per-glycan composition distribution and core-1:core-2 ratio for Thr143, derived from glycoproteomic analysis of peptide T[143]EVFT[147]PVVP following CT and IMPa digestion of untreated OPN.**

| Glycan type | Mass (Da) | Fraction of T143 |
|---|---|---|
| N1 | 203.0794 | 0.00 |
| N1H1 | 365.1322 | 0.12 |
| N2H1 | 568.2116 | 0.00 |
| N2H2 | 730.2644 | 0.08 |
| N1H1A1 | 656.2276 | 0.13 |
| N2H1A1 | 859.307 | 0.00 |
| N1H1A2 | 947.323 | 0.30 |
| N2H2A1 | 1021.3598 | 0.23 |
| N2H2A2 | 1312.4552 | 0.14 |
| Core-1 | | 0.55 |
| Core-2 | | 0.45 |

### References

(1) Marty, M. T.; Baldwin, A. J.; Marklund, E. G.; Hochberg, G. K. A.; Benesch, J. L. P.; Robinson, C. V. Bayesian Deconvolution of Mass and Ion Mobility Spectra: From Binary Interactions to Polydisperse Ensembles. *Anal. Chem.* **2015**, *87* (8), 4370–4376. https://doi.org/10.1021/acs.analchem.5b00140

(2) Schachner, L. F.; Mullen, C.; Phung, W.; Hinkle, J. D.; Beardsley, M. I.; Bentley, T.; Day, P.; Tsai, C.; Sukumaran, S.; Baginski, T.; DiCara, D.; Agard, N. J.; Masureel, M.; Gober, J.; ElSohly, A. M.; Melani, R.; Syka, J. E. P.; Huguet, R.; Marty, M. T.; Sandoval, W. Exposing the Molecular Heterogeneity of Glycosylated Biotherapeutics. *Nat. Commun.* **2024**, *15* (1), 3259. https://doi.org/10.1038/s41467-024-47693-8

(3) Riley, N. M.; Bertozzi, C. R. Deciphering O-Glycoprotease Substrate Preferences with O-Pair Search. *Mol. Omics* **2022**, *18* (10), 908–922. https://doi.org/10.1039/D2MO00244B

(4) Riley, N. M.; Malaker, S. A.; Driessen, M. D.; Bertozzi, C. R. Optimal Dissociation Methods Differ for N- and O-Glycopeptides. *J. Proteome Res.* **2020**, *19* (8), 3286–3301. https://doi.org/10.1021/acs.jproteome.0c00218

(5) Polasky, D. A.; Yu, F.; Teo, G. C.; Nesvizhskii, A. I. Fast and Comprehensive N- and O-Glycoproteomics Analysis with MSFragger-Glyco. *Nat. Methods* **2020**, *17* (11), 1125–1132. https://doi.org/10.1038/s41592-020-0967-9

(6) Lu, L.; Riley, N. M.; Shortreed, M. R.; Bertozzi, C. R.; Smith, L. M. O-Pair Search with MetaMorpheus for O-Glycopeptide Characterization. *Nat. Methods* **2020**, *17* (11), 1133–1138. https://doi.org/10.1038/s41592-020-00985-5

(7) Polasky, D. A.; Lu, L.; Yu, F.; Li, K.; Shortreed, M. R.; Smith, L. M.; Nesvizhskii, A. I. Quantitative Proteome-Wide O-Glycoproteomics Analysis with FragPipe. *Anal. Bioanal. Chem.* **2025**, *417* (5), 921–930. https://doi.org/10.1007/s00216-024-05382-x

(8) Ferries, S.; Perkins, S.; Brownridge, P. J.; Campbell, A.; Eyers, P. A.; Jones, A. R.; Eyers, C. E. Evaluation of Parameters for Confident Phosphorylation Site Localization Using an Orbitrap Fusion Tribrid Mass Spectrometer. *J. Proteome Res.* **2017**, *16* (9), 3448–3459. https://doi.org/10.1021/acs.jproteome.7b00337

(9) Daly, L. A.; Brownridge, P. J.; Batie, M.; Rocha, S.; Sée, V.; Eyers, C. E. Oxygen-Dependent Changes in Binding Partners and Post-Translational Modifications Regulate the Abundance and Activity of HIF-1α/2α. *Sci. Signal.* **2021**, *14* (692), eabf6685. https://doi.org/10.1126/scisignal.abf6685

(10) Janek, K.; Wenschuh, H.; Bienert, M.; Krause, E. Phosphopeptide Analysis by Positive and Negative Ion Matrix-Assisted Laser Desorption/Ionization Mass Spectrometry. *Rapid Commun. Mass Spectrom. RCM* **2001**, *15* (17), 1593–1599. https://doi.org/10.1002/rcm.417

(11) Daly, L. A.; Byrne, D. P.; Perkins, S.; Brownridge, P. J.; McDonnell, E.; Jones, A. R.; Eyers, P. A.; Eyers, C. E. Custom Workflow for the Confident Identification of Sulfotyrosine-Containing Peptides and Their Discrimination from Phosphopeptides. *J. Proteome Res.* **2023**, *22* (12), 3754–3772. https://doi.org/10.1021/acs.jproteome.3c00425

(12) Christensen, B.; Nielsen, M. S.; Haselmann, K. F.; Petersen, T. E.; Sørensen, E. S. Post-Translationally Modified Residues of Native Human Osteopontin Are Located in Clusters: Identification of 36 Phosphorylation and Five O-Glycosylation Sites and Their Biological Implications. *Biochem. J.* **2005**, *390* (1), 285–292. https://doi.org/10.1042/BJ20050341

(13) Froehlich, J. W.; Chu, C. S.; Tang, N.; Waddell, K.; Grimm, R.; Lebrilla, C. B. Label-Free Liquid Chromatography–Tandem Mass Spectrometry Analysis with Automated Phosphopeptide Enrichment Reveals Dynamic Human Milk Protein Phosphorylation during Lactation. *Anal. Biochem.* **2011**, *408* (1), 136–146. https://doi.org/10.1016/j.ab.2010.08.031

(14) Christensen, B.; Petersen, T. E.; Sørensen, E. S. Post-Translational Modification and Proteolytic Processing of Urinary Osteopontin. *Biochem. J.* **2008**, *411* (1), 53–61. https://doi.org/10.1042/BJ20071021

(15) Kariya, Y.; Kanno, M.; Matsumoto-Morita, K.; Konno, M.; Yamaguchi, Y.; Hashimoto, Y. Osteopontin O-Glycosylation Contributes to Its Phosphorylation and Cell-Adhesion Properties. *Biochem. J.* **2014**, *463* (1), 93–102. https://doi.org/10.1042/BJ20140060

(16) Li, H.; Shen, H.; Yan, G.; Zhang, Y.; Liu, M.; Fang, P.; Yu, H.; Yang, P. Site-Specific Structural Characterization of O-Glycosylation and Identification of Phosphorylation Sites of Recombinant Osteopontin. *Biochim. Biophys. Acta BBA - Proteins Proteomics* **2015**, *1854* (6), 581–591. https://doi.org/10.1016/j.bbapap.2014.09.025

(17) Mateos, B.; Holzinger, J.; Conrad-Billroth, C.; Platzer, G.; Żerko, S.; Sealey-Cardona, M.; Anrather, D.; Koźmiński, W.; Konrat, R. Hyperphosphorylation of Human Osteopontin and Its Impact on Structural Dynamics and Molecular Recognition. *Biochemistry* **2021**, *60* (17), 1347–1355. https://doi.org/10.1021/acs.biochem.1c00050

(18) Huang, H.; Arighi, C. N.; Ross, K. E.; Ren, J.; Li, G.; Chen, S.-C.; Wang, Q.; Cowart, J.; Vijay-Shanker, K.; Wu, C. H. iPTMnet: An Integrated Resource for Protein Post-Translational Modification Network Discovery. *Nucleic Acids Res.* **2018**, *46* (D1), D542–D550. https://doi.org/10.1093/nar/gkx1104
